# Osmotic stress response in BetP: How lipids and K^+^ team up to overcome downregulation

**DOI:** 10.1101/2022.06.02.493408

**Authors:** Veronika Heinz, Günnur Güler, Vanessa Leone, M. Gregor Madej, Stanislav Maksimov, Rebecca M. Gärtner, Olga Rudi, Farzad Hamdi, Panagiotis L. Kastritis, Werner Mäntele, Reinhard Krämer, Lucy R. Forrest, Camilo Perez, Christine Ziegler

## Abstract

The trimeric betaine symporter BetP senses an osmotic upshift via its osmosensory C-terminal domain and responds rapidly with a steep increase in transport rate. Full activation requires both an elevated internal K^+^ concentration and a yet unknown membrane stimulus. Moreover, the molecular mechanisms of stress sensing and upregulation remain unclear. Here, we show that K^+^ binding to BetP *in vivo* is highly cooperative. Using X-ray crystallography, we identify four putative K^+^ interaction sites at the C-terminal domains of BetP. Single particle CryoEM on BetP reconstituted in amphipols (AMP-BetP), in the absence of K^+^, revealed that the three C-terminal domains are oriented symmetrically near the membrane surface, representing a new downregulated state. The presence of K^+^ resulted in asymmetric partial unfolding of the C-terminal domains, which was assigned as an intermediate between the downregulated state and the conformation observed in crystal structures. Attenuated total reflection Fourier transform infrared (ATR-FTIR) spectroscopy in 2D crystals of BetP reveal glutamate/aspartate and tyrosine responses to K^+^, in agreement with the identified K^+^ interaction sites, as well as specific unfolding events in the C-terminal domain upon activation. A rearrangement of the relative protomer orientations confers upregulation through key structural elements involved in the alternating access of BetP affecting sodium and betaine binding affinities. Although K^+^-specific regulation is unique to BetP we discuss unfolding/refolding of sensory domains as a unifying element in hyperosmotic stress response of osmoregulated transporters.

## Introduction

The sodium-coupled betaine symporter BetP from the soil bacterium *Corynebacterium glutamicum* is a prime example of a lipid-dependent osmotic stress regulated membrane transporter [1]. BetP transports betaine with high specificity, which under hyperosmotic conditions is accumulated in molar amounts against the concentration gradient into the cytoplasm [2]. The transporter exploits the inward directed sodium gradient across the membrane to energize betaine transport in a stoichiometry of 2 Na^+^:1 betaine [3]. It is assumed that BetP senses changes of the physical state of the membrane via its C-terminal domain [1, 4, 5]. Biochemical data suggested that the sensory domain is attached at the membrane surface when BetP is downregulated in the absence of stress [6, 7] and that these specific lipid interactions are released when BetP perceives a stress signal via the membrane [8, 9]. Activation can be partially achieved by increasing K^+^ concentrations in proteoliposomes [4, 10], but notably, K^+^ alone - i.e., in the absence of the membrane stimulus - is not able to fully activate BetP [7]. *Vice versa*, full transport activity cannot be reached by the membrane stimulus alone [5, 7]. The biochemical properties of the C-terminal domain(s) have been studied extensively during the past decades and it was shown that osmo-stress regulation of BetP is strongly dependent on the helical folding of this osmosensory domain [7, 11]. Transport regulation triggered by hyperosmotic stress was seen in other transporters such as OpuA, a type I ABC importer, which was shown to be gated by ionic strength and inhibited by the second messenger cyclic-di-AMP [12, 13]. The specific requirement of potassium ions during activation and the characteristic helical C-terminal domain was, however, only observed in BetP [7]. In several crystal structures [2, 14–17] BetP appears as a trimer with each protomer in a different conformational state and each osmosensor exhibiting a different degree of helical folding (**Fig. S1**). Consistently, only one of the C-terminal domains, the one involved in crystal contacts, formed a straight alpha-helix. This domain was maximally resolved to Arg586, exhibiting two segments (HC1: Glu552 – Arg568 and HC2: Glu570 – Arg595) separated by an unwound region (**Fig. S1,** chain A). The other two C-terminal domains were resolved in most structures at most to Arg568 and at least to Glu552, respectively. Aside from several arginine residues in HC1 and HC2, Tyr550 in the short linker loop between TM12 and HC1, and Glu572 in HC2 are essential for osmosensing [7]. It was suggested that having positively charged residues in HC1 and HC2 is key to the lipid-dependent down-regulation in BetP. The membrane of *C. glutamicum* consists entirely of negatively charged lipids, with phosphatidylglycerol (PG) and cardiolipin (CL) together contributing nearly 70%. The osmotic activation threshold, directly dependent on the amount of negatively charged lipids in the membrane, shifts to lower osmolalities when BetP is expressed in *E. coli*, which comprises ~70% neutral lipids. Although BetP retains up to eight PG lipids bound mainly in the hydrophobic cavity in the trimer centre none of the C-terminal domains in crystal structures adopts a conformation approaching the membrane; in fact, all three C-terminal helices are pointing around 30° away from the membrane plane. To date, it has not been possible to assign a specific activation state to the crystal structures, and consequently neither a molecular mechanism for stress sensing nor activation could be deduced for BetP.

Here, we present X-ray and Cryo-EM BetP structures in complex with K^+^, which allow us to unambiguously assign down-regulated and activated conformations for BetP. By using a combination of functional, structural, spectroscopic, and computational studies, we pinpoint the pivotal role of lipids and K^+^ in downregulation and identify unfolding-refolding of the osmosensory C-terminal domain as a key element to overcome downregulation in response to elevated K^+^ concentrations.

## Results

### K^+^ binds to multiple sites in BetP

Regulation of transport activity of BetP as a function of the luminal K^+^ concentration was measured under both *in vivo* (*C. glutamicum* cells) and *in vitro* conditions (*E. coli* lipid proteoliposomes) and found to be highly cooperative (**Fig. 1**). The *in vivo* analysis is limited to *C. glutamicum* cells, since only this organism can be grown under extremely low internal potassium concentrations, which is an experimental prerequisite for this investigation [18]. Hill coefficients for K^+^ dependent activation were directly deduced from Fig.1 resulting in n = 4.9 ± 0.2 and 5.2 ± 0.2 in *C. glutamicum* cells and proteoliposomes, respectively, suggesting the presence of cooperative K^+^ binding in BetP during activation. The corresponding half-activation constants for K^+^ were 220 ± 14 mM and 242 ± 15 mM in *C. glutamicum* cells and proteoliposomes, respectively. We therefore considered the *K*d as a threshold of activation for subsequent structural and spectroscopic studies. To further characterize the nature of cooperative K^+^ binding, we co-crystallized BetP with the K^+^ mimic Rb^+^.

**Figure 1.**
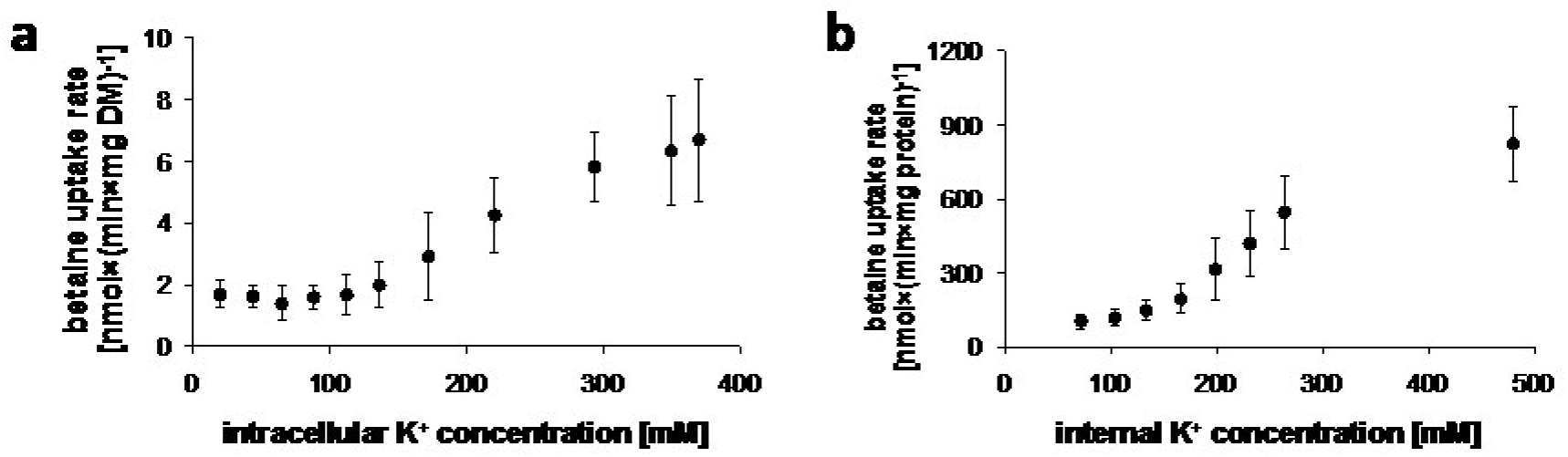
Effect of internal K^+^ concentration on BetP activity. **a.** [^14^C]-Betaine uptake rate of BetP expressed in *C. glutamicum* cells. Cytoplasmic K^+^ concentration was gradually increased by K^+^ uptake into K^+^ depleted cells. **b.** [^14^C]-Betaine uptake rate of BetP reconstituted in liposomes. The internal K^+^ concentration was increased by controlled shrinkage of the vesicles in response to addition of hypertonic buffer. All measurements were carried out in triplicate and error bars indicate standard deviation (s.d.).

### BetP co-crystallization with Rb^+^

Activation of BetP can also be achieved by Rb^+^ and Cs^+^ [10]. Rb^+^ has a comparable ionic radius (*r*) to that of K^+^ (*r*^Rb+^ = 166 pm; *r*^K+^ = 152 pm), so a similar coordination of Rb^+^ and K^+^ ions in BetP can be assumed. Therefore, we co-crystallized BetP with the K^+^ analogue Rb^+^ [19, 20] to trace protein-K^+^ interaction-sites via anomalous Rb^+^ scattering. According to our *in vivo* cooperativity measurements, we used 300 mM K^+^ (Rb^+^) for co-crystallization, which would be above the *K*_d_ in the presence of 1 mM betaine. We solved the structure of the Rb^+^-complexed BetP to a resolution of 3.4 Å (**Fig. 2a**). All three protomers adopt an inward-facing open C_i_ conformation similar to a structure reported previously [2]. Chain A contains a betaine molecule bound close to the S1 site that is coordinated through cation-π interactions with Trp377 similar to previous structures [15]. The C-terminal domain of chain A is resolved as a straight helix as far as residue Arg584 (**Fig. 2a**) and projects towards chain C, which harbours a citrate buffer molecule in the cytoplasmic pathway. The C-terminal domain of chain C is resolved up to residue Arg568 and pointing towards chain B, which adopts an apo state, i.e., neither substrate nor buffer molecule occupies the intracellular pathway.

**Figure 2.**
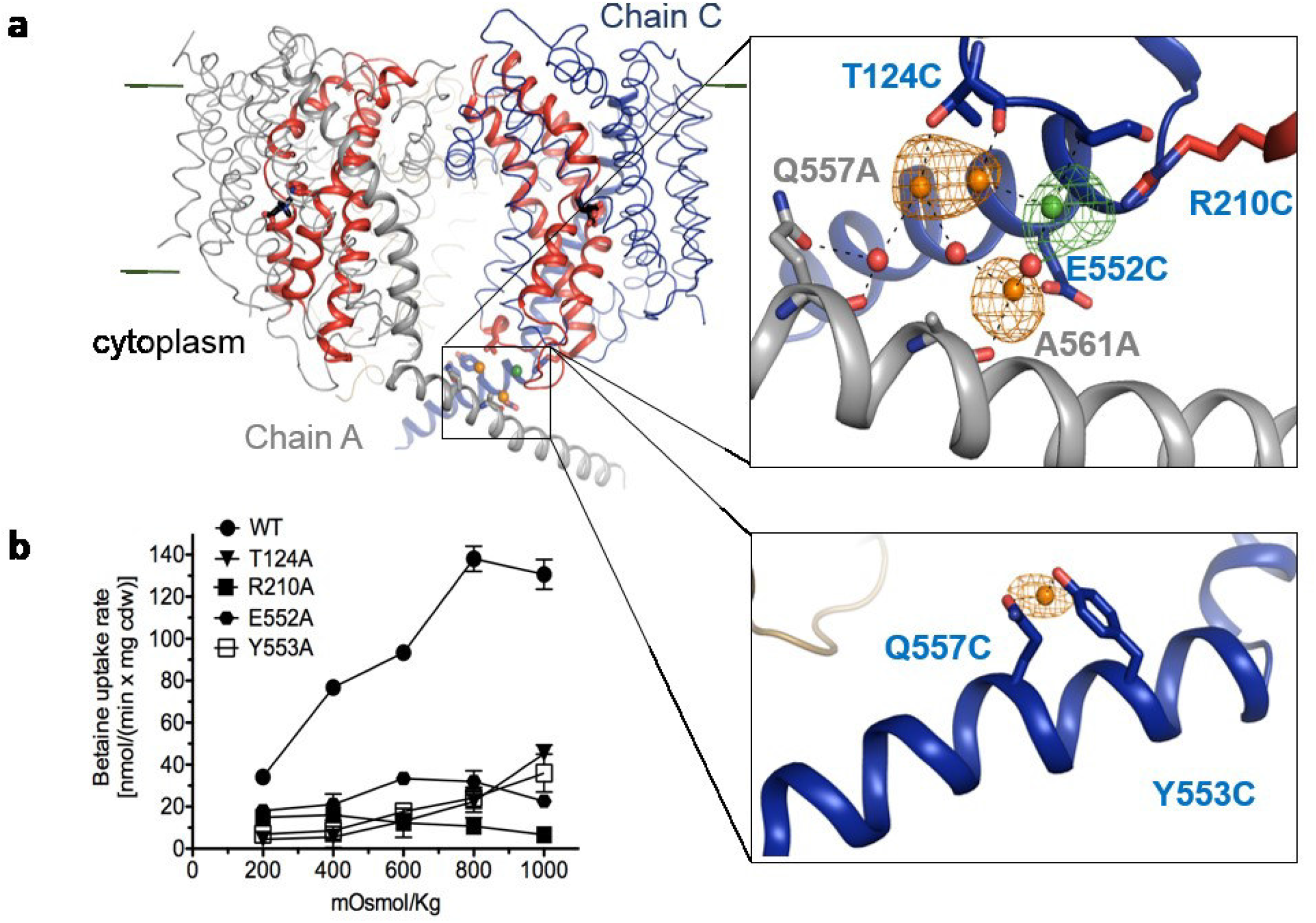
The role of K^+^ in the regulation of BetP. **a.** Rb^+^ binding sites in BetP. Rb^+^ ions (orange) and their interaction networks are shown with the anomalous-difference Fourier map (orange) contoured at 4.0σ. The *Fo-Fc* omit map (green) of the coordinating Cl^−^ ion (green) is contoured at 3.0σ. Water molecules are shown as red spheres. Betaine and citrate are shown in black sticks. **b.** Functional characterization of Rb^+^ interaction-sites. Osmotic activation profile of BetP WT expressed in *E. coliMKH13* compared to those of single alanine mutants of residues involved in the formation of Rb^+^ interaction sites. All measurements were carried out in triplicate. Error bars indicate s.d.

Anomalous difference Fourier maps were calculated using the CCP4 program FFT [21] and allowed for an unambiguous assignment of Rb^+^ in the crystal structure. Four cytoplasmic Rb^+^ ions are observed (**Fig. 2a insets**). Three Rb^+^ ions are localized at the interface of the crossing C-terminal domains from chains A and C with the first one coordinated by the carboxyl group of Glu552 from the C-terminal domain of chain C, the main chain carbonyl of Ala561 from the C-terminal domain of chain A, and a hydrated Cl^-^ ion close to the side chain of Arg210 from chain C. A second Rb^+^ ion is coordinated by the hydroxyl group of Thr124 from chain C, a water molecule hydrogen-bonding to the main chain carbonyl, and the side chain of Gln557 from the C-terminal domain of chain A, and the third Rb^+^ ion is coordinated by the carbonyl of Thr124 and the hydrated Cl^-^ ion. A fourth Rb^+^ ion is coordinated only by residues located in the C-terminal domain of chain C, including the side chain of Gln557 and the hydroxyl group of Tyr553 (**Fig. 2a insets**). To study the contribution of each site to the osmo-regulation activity profile, we performed alanine replacement of the cytoplasmic side chains involved in the Rb^+^ interaction sites (**Fig. 2b**). Alanine mutants of residues Thr124, Arg210, Glu552 and Tyr553 exhibit severely altered responses to osmotic stress. While R210A and E552A are no longer regulated, T124A and Y553A show a slight increase in activity at high osmolalities. The alanine mutant of Gln557 (**Fig. 3a**) retains the ability to respond to osmotic stress although with a significant shift to higher osmolalities resulting at a reduced transport rate. In summary, almost all mutants of the residues participating in the cytoplasmic Rb^+^ interaction sites showed impaired osmo-sensing and/or reaction properties compared to WT BetP. The mutagenesis study confirms the Rb^+^-interacting residues are essential for activation and that they are located mainly in HC1. Again, the Rb^+^ bound structure shows only very slight deviations from structures determined in the absence of Rb^+^ / K^+^ (e.g., PDB ID 4C7R) suggesting that the ensembles of C-terminal domains, especially the crossing of C-terminal domains of chain A and chain C in the trimer already represent an active conformation with K^+^ interaction sites pre-formed. We investigated the C-terminal domain inter-protomeric interactions by measuring the formation of a crosslink between the C-terminal domain and L2 of adjacent protomers mimicking the interaction between Arg558 and Asp131 observed in crystal structures (**Fig. S1**), as a function of osmotic stress (**Fig. 3b**). First, residues Arg565 and Ile130 were replaced with cysteine; this mutant was expressed in *E. coli* cells and the osmotic profile was measured to test whether its activity could still be osmo-regulated. The BetP-R565C/I130C mutant was also regulated by changes in osmotic stress, although the initial activity measured at 200 mOsmol/kg was ~2.5-fold higher than WT, and the overall increase of activity was only 1.5-fold, compared to ~4-fold for the fully regulated WT BetP (**Fig. 3b**). We compared WT and the cross-linked BetP-I130C/R565C using SDS-PAGE and immunoblotting against the N-terminal strepII tag after the addition of the homo-bifunctional cross-linker o-PDM. There is a significant amount of dimeric cross-linked BetP-I130C/R565C at low osmolalities (**Fig. 3b,** blue square), with limited amounts of trimeric protein, whereas at high osmolality the cross-linked protein exists in both dimeric and trimeric forms (**Fig. 3b,** green square). These results indicate that the interaction between the C-terminal domain of one protomer and L2 of an adjacent protomer at this position is characteristic for an activated state. Therefore, we can conclude that the C-terminal conformations observed in crystal structures with and without K^+^ represent an active state. There might be several reasons that a down-regulated state cannot be adopted in crystal structures, such as missing lipids in the detergent micelle or the altered activation profile of the crystallization mutant. However, ionic strength and crowding effects during crystallization could simply prevent downregulated conditions. To test this hypothesis, we performed a CryoEM single particle analysis on WT BetP.

**Figure 3.**
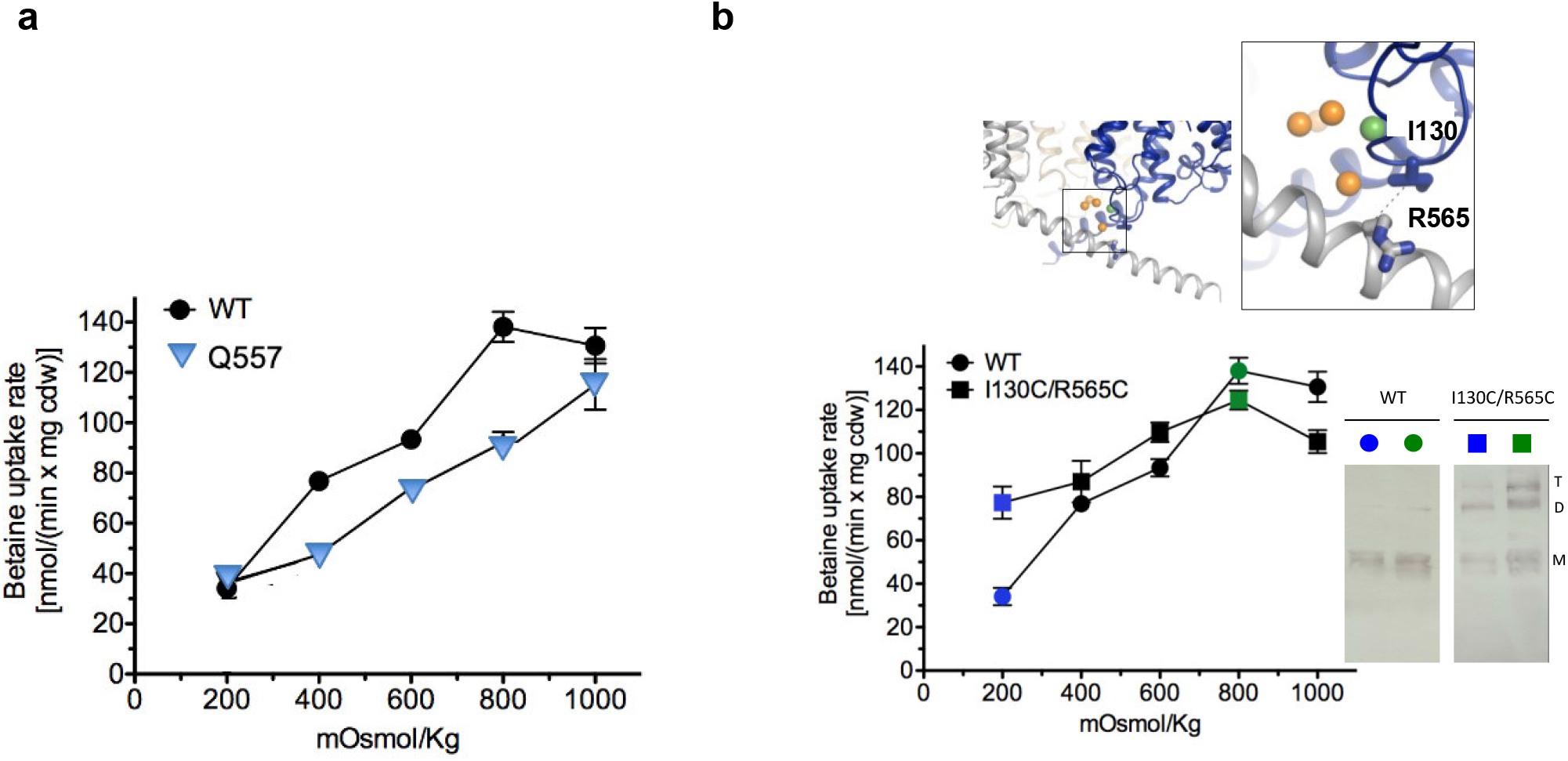
The active state of BetP in crystal structures. **a.** Osmotic activation profile of BetP WT expressed in *E. coli MKH13* compared to that of single alanine mutants of Gln557 coordinating Rb^+^. All measurements were carried out in triplicate. Error bars indicate s.d. **b.** Osmotic activation profile of BetP WT and the double cysteine mutant I130C/R565C expressed in *E. coli MKH13* cells and western blot of BetP WT and I130C/R565C from membrane vesicles of *E. coli MKH13* cells extracted after incubation in iso-osmotic conditions (blue) or hyper-osmotic conditions (green), in the presence of the homobifunctional cross-linker o-PDM (o-phenylenedimaleimide). T, trimer; D, dimer; M, monomer.

### The downregulated state of amphipol-reconstituted BetP by CryoEM

Considering the strong effect of negatively charged lipids on BetP activation we reconstituted WT BetP into amphipol A8-35, which due to its carboxyl-groups provides a negatively charged belt around the BetP trimer (**Fig. S2a**). In comparison to BetP reconstituted in *E. coli* proteoliposomes and POPG-MSP nanodiscs, respectively, amphipol-reconstituted BetP (AMP-BetP) showed comparable sodium-dependent betaine binding (**Fig. S2b, Table S1**). Moreover, binding was responsive to K^+^ concentrations (**Fig. S2c, Table S1**) with a similar increase in the Hill coefficient to ~2 as observed for BetP reconstituted in proteoliposomes at elevated K^+^ concentrations [22]. CryoEM micrographs of AMP-BetP were collected in the absence of K^+^ (**Table S2, Fig S3**). Two distinct populations were identified. The larger class showed all three C-terminal domains resolved to comparable length and arranged symmetrically and parallel to the membrane plane (**Fig. S3 a, d**). The second, significantly smaller 3D class resulted in a 3D volume, in which the C-terminal domains were resolved to Arg568, e.g., only HC1 was folded (**Fig. S3a**). We also investigated the crystallization mutant of BetP (BetP^cryst^ - ΔN29 E_3_ 44-46 A_3_) reconstituted in amphipol by CryoEM. This mutant displayed a higher conformational heterogeneity in which the dominant class adopted an asymmetric conformation with HC1 folded and HC2 unfolded to different degrees, as in the second class of WT BetP (**Fig. S3a, Fig. S4**). The C-terminal orientations in the CryoEM coulomb potential maps of BetP and BetP^cryst^ were more or less like each other but were distinct from the orientation observed in crystal structures (**Fig. S1**).

The C3 map of the symmetrical class of WT BetP reconstructed to 3.7 Å (**Fig. 4a**) was modeled (**Fig. 4b, Table S4**). All three C-terminal domains in the CryoEM structure are resolved to Ala580 and form a straight helix. The cytoplasmic loops L2 and L8 form two short, anti-parallel beta-strands between residues Ile125 – Leu127 (L2) and Gly391 – Ile394 (L8) (**Fig. S5)**. In comparison to crystal structures, we observe a rigid-body rotation of the C-terminal helix around Asp547 by 50° towards the outer rim of the trimer (**Fig. S6a**). Simultaneously, the C-terminal domain is tilted upwards by 10° towards the membrane plane (**Fig. S6b**). In this orientation it points towards loop 6 (L6) and TM1 of the adjacent protomer (**Fig. S6c, d**). We analysed whether this orientation leads to persistent contacts during μs-scale molecular dynamics (MD) simulations of the BetP trimer inserted into a POPG lipid membrane. In the simulation initiated with the CryoEM orientation of the protomers, the C-terminal domain forms extensive contacts with the cytoplasmic loops L2, L6, L8 and L10 (**Fig. 76 a, b**). These are in stark contrast with the contacts observed in a simulation initiated with an X-ray structure, where only a handful of the contacts with L2 are similar (**Fig. S7 a, b**). Notably, in the CryoEM structure, the protomer distances within the trimer differ by 2.6 Å with respect to the threefold symmetry axis at the cytoplasmic side in comparison to the X-ray structure (e.g., between C_α_ of Lys121 in chain A, B and C), resulting in a narrower hydrophobic cavity. TM1, TM6 and TM11 are oriented parallel to the membrane normal, while they are tilted away from the membrane normal in X-ray structures (**Fig. S6b)**. During the MD simulations of the trimer in a POPG environment for both X-ray and CryoEM structures, the interprotomer distances evolve during the simulations (**Fig. S7c, d**), suggesting that the lipid bilayer is less constraining than the amphipol conditions; nevertheless, the trimer remains significantly more constrained and symmetric in the CryoEM-derived conformation than in the X-ray structure.

**Figure 4.**
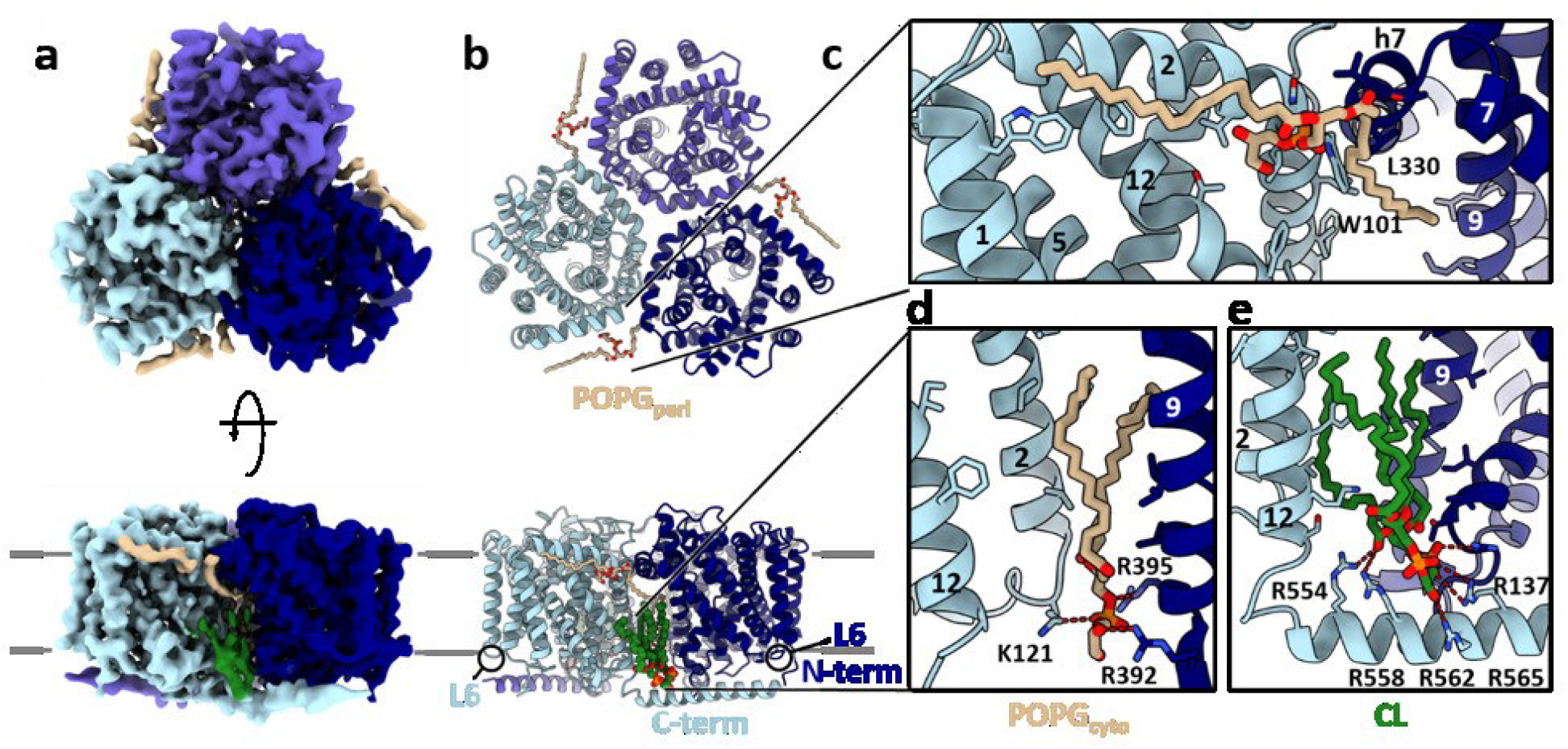
Symmetric cryo-EM structure of BetP in the absence of K^+^. **a.** Cryo-EM density map of BetP in C3 symmetry at a resolution of 3.7Å according to the FSC = 0.143 criterion. The BetP trimer (protomer A: light blue, protomer B: violet blue, protomer C: navy blue) is depicted in a top- and side-view. Lipid densities located at the hydrophobic cleft at the trimer interface are shown in tan (POPG) or green (CL), respectively. The membrane plane is indicated by grey lines. **b.** An atomic model was built into the density map shown in a. Intra-protomeric contacts involve the N- and C-terminal domains and L6 as well as the lipid molecules. Color coding as in a. **c.** Coordination of the POPG^peri^ molecule with TM1, TM2 and TM12 in protomer A and h7 and TM9 in protomer C. **d.** Coordination of POPG^cyto^ with TM2 and TM12 in protomer A and TM9 in protomer C. **e.** Coordination of CL with TM2, TM12 and the C-terminal domain of protomer A and TM9 and L2 in protomer C.

In summary, we assign the symmetrical trimeric state observed for AMP-BetP as a down-regulated conformation, noting that the wildtype AMP-BetP contains a substantial fraction (24.1%) of trimers that are, according to this definition, not in a down-regulated state (**Fig. S3a**). We assign this population to an active state, similar to other asymmetrical trimeric states, in which C-terminal domains differ in degree of helical folding and/or orientation. Unlike the wild-type protein, BetP^cryst^ does not exhibit any population of the fully symmetrical downregulated state when reconstituted in amphipol, although the HC1 helix of the C-terminal domains lies parallel to the membrane, unlike in the crystal structures (**Fig. S4b**). _As BetP_^cryst^ _is_ partly N-terminal truncated we suggest that the helical stabilization of HC2 required for the fully-symmetric state is normally facilitated by N-terminal interactions. Therefore, BetP^cryst^ can never be entirely down-regulated, consistent with the altered activation profile of this and other N-terminally truncated mutants [1, 14].

### Lipid binding to AMP-BetP

Elongated densities identified as lipids are located in the hydrophobic clefts between the protomers and at the trimer perimeter (**Fig. 4**). The lipid densities were assigned as POPG and cardiolipin (CL), respectively. We also detected a similar density for cardiolipin in AMP-BetP^cryst^ (**Fig. S4b**). A periplasmic POPG (POPG^peri^) is resolved close to the amphipatic helix h7 at a trimerization contact between Trp101 in TM2 and Leu330 in h7 [23], while a cytoplasmic POPG (POPG^cyto^) is found in the hydrophobic cleft coordinated by Lys121 (L2) and by Arg392 and Arg395 (L8) of the counterclockwise adjacent protomer (**Fig. 4c, d**). The cardiolipin fatty acid chains protrude into the cleft formed between protomers (**Fig. 4b, e**), while the cardiolipin phosphate group is nestled into an arginine-binding pocket formed by Arg554, Arg558, Arg562 and Arg565 in the C-terminal domain and by Arg137 in L2. In addition, cardiolipin interacts with POPG^cyto^ and with hydrophobic residues in TM2/TM12/loop 4 (L4) and TM3/TM9/L2 of the protomers flanking the cleft. Being positioned between the cardiolipin molecule and L2, the C-terminal domain of the clockwise adjacent protomer maintains several ionic interactions starting from Arg554 to Ala580 with L2, which is consistent with peptide binding studies [7] (**Fig. 4e**).

### Inward-facing conformation in down-regulated AMP-BetP

All three protomers adopt an apo-inward facing state (C_i_^down^), which compared to those observed in crystal structures (C_i_^cryst^), exhibits differences in TM3, TM7, h7 and the cytoplasmic loops L2, L4, L6, and L8 (**Fig. 5a-c**). Differences in TM7 and h7 are correlated with an interaction between Lys300 in L6 and Glu572 of the C-terminal domain of the clockwise adjacent protomer (**Fig. S6d**). Specifically, L6 is stretched out away from the transporter core affecting the orientation of TM7 and h7. L6 and the cytoplasmic tip of TM7 were already identified as key players in conformational cycling in BetP, but also in other LeuT-fold transporters [24]. Mutation of K300A and G301A resulted in inactive BetP mutants, while BetP E572P lost transport regulation [5]. TM3 is separated by a conserved glycine stretch (G149-M150-G151-I152-G153) into two helical segments (TM3_a_ and TM3_b_). In AMP-BetP the cytoplasmic TM3_a_ exhibits a slightly compressed glycine stretch and Met150 points towards the Trp prism interacting with Trp374 (**Fig. 5c**). This rotamer conformation was not observed in Ci^cryst^, where the Met150 thioether side chain points towards the hydrophobic cleft formed by TM3 and TM7 interacting with the fatty acid chain of a POPG molecule.

**Figure 5.**
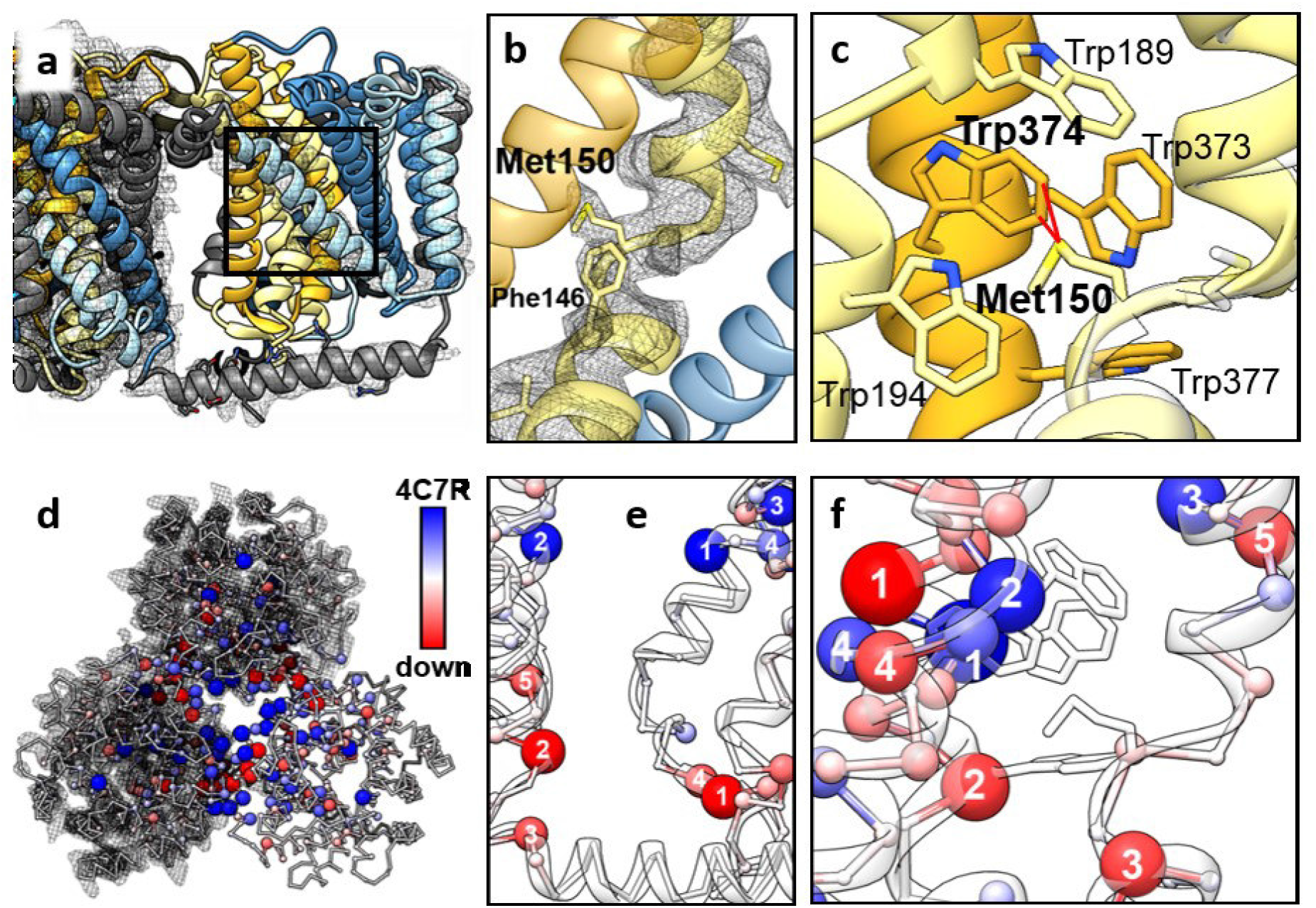
Structural analysis of BetP in the absence of K^+^. **a.** Superposition of the atomic ribbon model with the cryo-EM density map, illustrating the tight interactions of the C-terminal domain with L2 and L6. A black box highlights the area shown in b and c. **b.** Close up of the glycine-stretch in TM3 (khaki). The atomic model is shown fit into the symmetric cryo-EM density map (black mesh). The neighboring TMs 9 (gold) and 10 (steel blue) are displayed as transparent ribbons. **c.** Superposition of the atomic model (gold/khaki) with crystal structure 4C7R (white). Met150 exhibits a significantly different orientation. In the absence of K^+^, the side chain is oriented towards the Trp box (Trp373, Trp374 and Trp377) in TM8 (gold ribbon, black mesh). **d.** Protein interaction network analyses were performed for the symmetric C_i_^down^ state and C_i_^cryst^ (pdb-ID 4C7R), each without ligands. The difference in global interaction of a specific residue was mapped on the atomic model of C_i_^down^ (white ribbon). Blue spheres indicate an increase in interaction in C_i_^cryst^, while red spheres indicated an increase in C_i_^down^. The sphere size corresponds to the number of interactions, i.e., large, red spheres represent residues with a higher interaction score under downregulated conditions. Most global interactors was identified at the trimer interface and in the bundle helices TM3, TM4, TM8 and TM9. **e.** A zoom on the contact region of protomers B and A reveals an altered interaction network at the regulatory C-terminal domain. For the C_i_^down^ state (red), residues Arg126.A (1), Ser545.B (2), Tyr550.B (3), Ile125.A (4) and Val541.B (5) are identified as major interactors, while in C_i_^cryst^ (blue) Phe112.A (1) and Phe112.B (2) as well as His192.A (3) and Phe110.A (4) contribute to the inter-protomer contact. **f.** A zoom on the Trp box in protomer C reveals an altered interaction profile in the substrate binding pocket: Trp373 (1), Trp189 (2), Tyr157 (3) and His192 (4) were identified as the key residues in C_i_^cryst^ (blue). Trp371 (1), Trp377 (2), Phe146 (3), Leu191 (4) and Phe156 (5) exhibit the highest interaction score in C_i_^down^ (red).

To assess the possibility that these two opposite conformations of Met150 are accessible just by a simple side chain reorientation, we analysed multiple μs-scale MD simulations of the AMP-BetP trimers initiated with the orientation of Met150 observed in a crystal structure in the presence of 100 mM and 300 mM K^+^. According to a metric of proximity (see Methods section) between Met150 and either Ala313, Ala408 and Trp412 (in TM3 and TM7, X-ray conformation) or Trp373, Trp374 and Trp377 in the Trp prism (EM conformation), the methionine almost exclusively points away from the Trp prism (**Fig. S8a**), although very rare but reversible transitions toward the Trp prism (e.g., CN(EM) values > 2) are observed. This is notable, given the use of a non-polarizable force field in which methionine-aromatic interactions have not been specifically optimized, and suggests that conformational rearrangements of Met150 in the down-regulated conformation are physiologically meaningful. We then investigated the possibility that the alternate conformation of Met150 depends on the orientation of the C-terminal domain. However, simulations carried out with the C-terminal helix oriented as in the X-ray structures did not cause a rearrangement of Met150 to interact with Trp374 (**Fig. S8b**). In summary, the simulations indicate that conformational rearrangements of Met150 observed in the CryoEM structure under down-regulated conditions are not related to the release of C-terminal domain interactions with cytoplasmic loops. Thus, it is possible that lipid interactions contribute to stabilising the unique side chain conformation of Met150 in the down-regulated AMP-BetP.

### CryoEM structure of AMP-BetP in the presence of K^+^

To investigate K^+^-induced structural changes of AMP-BetP, a CryoEM dataset was collected in the presence of 200 mM KCl (**Fig. S9a-c**, **Table S2**). The CryoEM coulomb potential map in C1 was reconstructed to 4.3 Å. AMP-BetP in the presence of K^+^ lacks C3 symmetry in the C-terminal domains, which all exhibit different lengths and orientations reminiscent of what was observed in crystal structures (**Fig. 6a**). Again, one of the C-terminal domains is helically folded until Ala576, while the C-terminal domains in protomers B and C are only resolved until Asn560 and Arg567, respectively with slightly different orientations to each other. We observed a low-resolution connecting density between the N-terminal domain and the C-terminal domain of the counterclockwise adjacent protomer (**Fig. S9d**). K^+^-AMP-BetP bridges between C_i_^down^ and C_i_^cryst^ adopting an intermediate, transient state C_i_^trans^. Differences are detected in L6, which is no longer interacting with the C-terminal domain. In turn the hairpin TM6 - TM7 is shifted towards the periplasmic side. Compared to protomer A (**Fig. 6c**) the C-terminal domain is folded helically until Arg565 still interacting with the cardiolipin molecule in protomers B and C (**Fig. 6d, e**). Moreover, the C-terminal domain in protomer C exhibits a different orientation more like that in crystal structures, which re-positions cardiolipin away from the membrane plane (**Fig. 6e**), which was not observed in BetP^cryst^ in the absence of K^+^ (**Fig. S4b**). Compared to C_i_^down^, C_i_^trans^ showed a conformation of the glycine stretch and Met150 like the one in C_i_^cryst^ confirming the MD simulation results that in the presence of K^+^, the orientation of Met150 away from the Trp prism is stabilized (**Fig. S8).** We therefore assign the inward-facing states observed in K^+^-AMP-BetP and in BetP crystal structures as pre-activated states.

**Figure 6.**
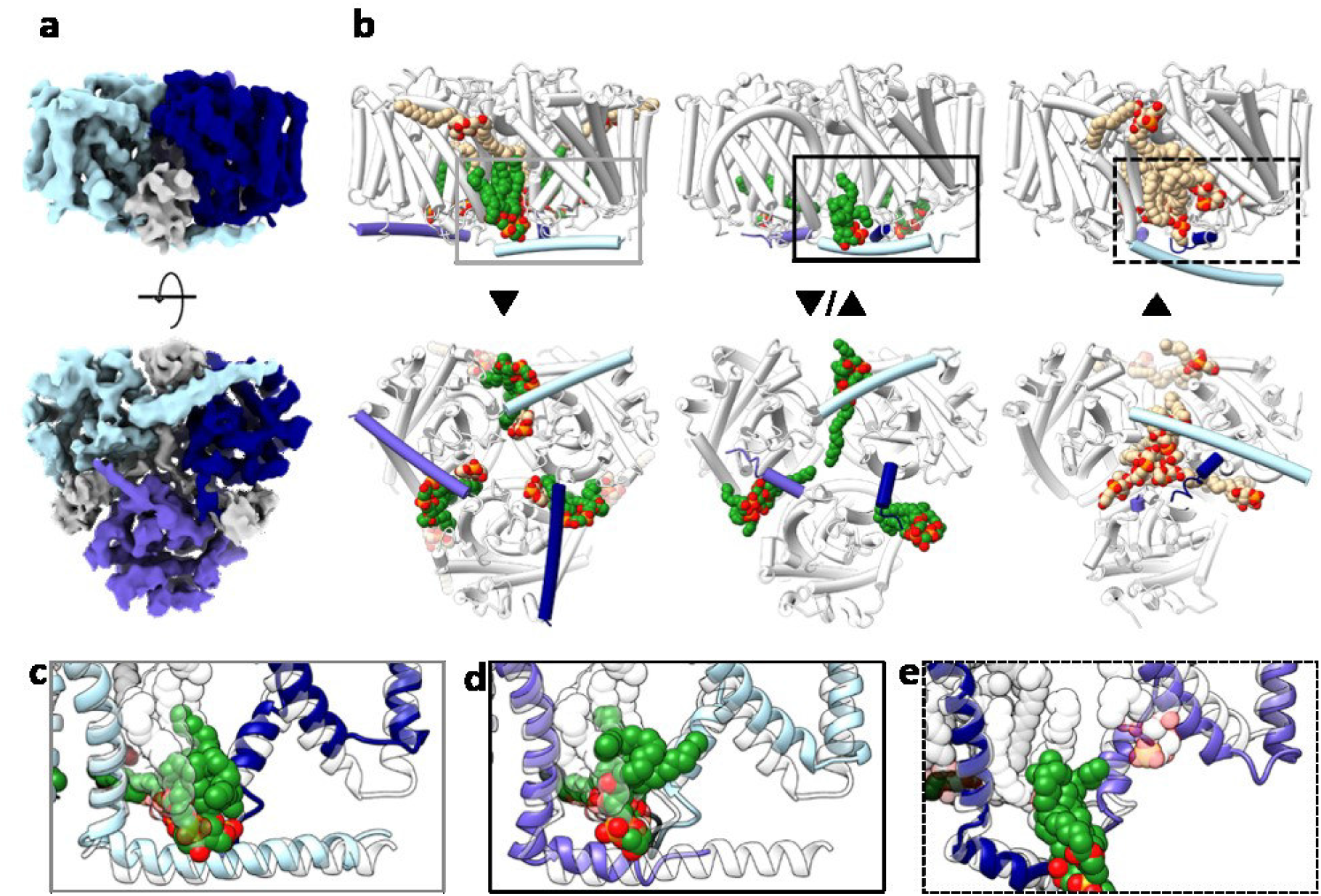
The effect of K^+^ on the structure of BetP. **a.** Asymmetric cryo-EM density map of BetP in the presence of 200 mM K^+^ at a resolution of 4.3 Å. The three protomers exhibit structural differences in length and orientation of the C-terminal domains: protomer A (light blue, fully extended), protomer B (violet blue, intermediate-length) and protomer C (navy blue, short). Lipid densities are depicted in grey. **b.** Structure and orientation of the C-terminal domains (color-coded according to a) of BetP in absence (C_i_^down^,**▼**) and presence (**▲**/**▼**) of K^+^, in comparison to crystal structure 4C7R (C_i_^cryst^,**▲**). Changes in the C-terminal domains are accompanied by altered lipid interactions with CL (green) and POPG (tan). **c.** Focus on the interaction of the C-terminal domain in protomer A (light blue) of BetP in the presence of K^+^ (**▼**/**▲**) with L2 and L6 of the adjacent protomer (navy blue, grey box in b), in comparison to C_i_^down^ (transparent). **d.** Focus on the interaction of the C-terminal domain in protomer B (violet blue) of BetP in the presence of K^+^ (**▼**/**▲**) with L2 and L6 of the adjacent protomer (light blue, black box in b), in comparison to C_i_^down^ (transparent). **e.** Focus on the interaction of the C-terminal domain in protomer C (navy blue) of BetP in the presence of K^+^ (**▼**/**▲**) with L2 and L6 of the adjacent protomer (navy blue, dashed box in b), in comparison to C_i_^cryst^ (transparent).

### Network-analysis of the down-regulated and transient states

Regulation in BetP is linked to the trimeric state. A network analysis was performed to identify global changes in interaction patterns in C_i_^cryst/trans^ and C_i_^down^ within the trimer (**Fig. S10**). Interactions were classified according to a metric that ranks the impact of a residue by the number of global interactions. We screened specifically for residues that changed their interaction pattern when switching from C_i_^down^ to C_i_^cryst^ (**Fig. 5d**). We detected changes in the intratrimeric contacts in residues located in the beta-strand in L2, which are highly interactive only in C_i_^down^, while residues located in helix h7 at the periplasmic side of the BetP trimer have increased interaction pattern only in C_i_^cryst^ (**Fig. 5e**), consistent with our observation of an altered protomer arrangement in downregulated AMP-BetP. With respect to substrate binding Trp371 and Trp377 are global interactors in C_i_^down^, while Trp373 and Trp189 have a higher interaction score in C_i_^cryst^ (**Fig. 5f**). Met150 shows equal impact on the global protein structure in both states, which we see in agreement to the importance of both alternating orientations as discussed before. An interaction network analysis including lipid molecules was performed to identify groups of global interactors with the C-terminal domain. In the C_i_^down^ state, two possible scenarios were identified. In the first scenario (**Fig. S11a**) the entire C-terminal domain forms a group with the transporter core of the counterclockwise adjacent protomer, while the cardiolipin lipid is grouped with the protomer providing the C-terminal domain. In the second scenario, cardiolipin is associated with the counterclockwise adjacent protomer, but the C-terminal domain is not. In fact, the C-terminal domain is only grouped until the very first segment (Arg558) with its own protomer (**Fig. S11b**). This pattern of partly grouping with the own protomer is maintained in the C_i_^cryst^ (**Fig. S11c, d**) suggesting a switch mechanism of C-terminal domain and lipids in BetP to stabilize the cytoplasmic part of the transporter core. Obviously, if the C-terminal domain is not engaged in intratrimeric interactions, it is not part of the group, which might cause a destabilization of the helical fold.

### K^+^ activation of lipid reconstituted BetP investigated by ATR-FTIR

The IR-absorbance spectra in the range 1800-1300 cm^-1^ were recorded from BetP in POPG 2D crystals at different concentrations of K^+^ (10 mM to 500 mM). For absorbance difference spectra, data were analysed after subtraction of the IR signal recorded in the absence of K^+^ (IR [10-500 mM K^+^]-IR [0 K^+^]) (**Fig. S12a**). The IR signal was further investigated by calculating the 2^nd^ derivative for peak assignment (**Fig. S12b-d)**. As vibration/stretching modes of individual residues and secondary structure elements often overlap in wavenumbers, spectra were taken in H_2_O and D_2_O, respectively, for unambiguous assignments of band position and band shift. This is necessary especially for the absorption of the H-O-H bending vibration at around 1643 cm^-1^ that superimposes with the amide modes (**Fig. S13a, b**). The deconvoluted bands were investigated for a possible assignment (**Fig. S12c, d**) to lipid C=O modes, to the C=O stretching modes of protonated Asp/Glu, to the Amide I region (i.e., to α-helices, unordered structures and loops), to the antisymmetric and symmetric stretching C-N mode of Arg residues, to the antisymmetric and symmetric stretching modes of deprotonated carboxylate groups for Asp/Glu, and to the C-C ring stretching vibrations of Trp residues and protonated/deprotonated Tyr residues, respectively. Multiple changes were observed in the 2^nd^ derivatives (**Fig. S12b-d**) and in the absorbance difference spectra (**Fig. S13**) from 10 – 200 mM (below the *K_d_* of K^+^) and from 250 – 500 mM K^+^ (above the *K*_d_ of K^+^). Analysis of the band shifts in the IR region of 1725 - 1700 cm^-1^, 1585 – 1565 cm^-1^ and 1390 – 1420 cm^-1^ allowed the assignment of distinct shifts in 1745, 1731, and 1715 cm^-1^, respectively, to changes in H-bonding of protonated Asp/Glu residues (**Fig. 7a**). Wavenumber shifts increased by ~ 6 – 8 cm^-1^ into saturation until 200 mM, followed by a steep decrease between 200 mM and 300 mM back to their initial peak position. Simultaneously, the IR signals originating from deprotonated Asp/Glu residues show the inverse spectral shift. Especially the negative absorbance between 1600 and 1550 cm^-1^ is an indicator of the population of deprotonated Asp/Glu residues. An increase in the population of stronger H-bonded protonated Asp/Glu side chains is only observed at K^+^ concentrations < 200 mM (below the *K*_d_ of K^+^) (**Fig. 1**), while above those concentrations the IR signal rapidly returns to its initial value. We suggest that Asp/Glu residues reporting changes from downregulated to active state might be transiently engaged in K^+^ binding as they return to an initial deprotonated state after full activation. We have identified potential K^+^-responsive Asp/Glu residues in the network analysis comparing C_i_^down^ and the activated C_i_^cryst/trans^ states. Asp470 (TM10), Asp543, and Asp547 (TM12) – all accessible from the cytoplasmic side - are strong interactors in the down-regulated state, while Asp131 exhibits a stronger interaction network in the activated state (**Fig. 7d**). Glu552, which is one of the K^+^ coordination sites identified in the Rb^+^ bound crystal structure and Glu572 have more interaction impact in the active state, Glu132 and Glu396 in L2 and L8 are more dominant in the downregulated state (**Fig. 7e**). Additional recording of the absorbance differences in D_2_O buffer confirms that absorbance differences at 1604 - 1587 cm^-1^ (**Fig. S13b**) can be assigned to the antisymmetric and symmetric stretching C-N mode of arginine, respectively, while absorbance differences at 1704 - 1692 and 1614 - 1611 cm^-1^ (**Fig. S12a**) are unambiguously assigned to the C=O mode and NH_2_ bending mode of asparagine, respectively (for band assignments see refs. [25–28]). These findings agree with our observation that the cardiolipin molecule remains bound to the Arg binding pocket and moves together with the C-terminal domain (**Fig. 6c-e**) therefore not contributing to the IR difference spectra. This observation would also agree with the fact that Arg210, which coordinates K^+^ indirectly via the chloride ion, is not affected in its conformation by K^+^ titration. In the difference spectra, the C-C ring mode of tyrosine, which is a sensitive protonation reporter group, also responds to a K^+^ increase (**Fig. 7b**). There is a distinctive upshift for the C-C mode of protonated Tyr (1516.5 cm^-1^) with a sudden breakdown after 200 mM K^+^, reminiscent of the upshift of protonated Asp/Glu residues. However, the C-C mode changes of deprotonated Tyr (1497 cm^-1^) respond differently to an increase in K^+^: the downshift from 10 - 100 mM K^+^ is followed by a lag phase between 100 mM - 200 mM K^+^. At 200 mM there is a steep bi-phasic increase, surpassing the initial wavenumber of 1496.8 cm^-1^ and reflecting the osmoregulation profile of BetP. Based also on our mutagenesis study (**Fig. 2b**) we assign Tyr553 as one of the key tyrosine residues to K^+^ activation (**Fig. 7f**). While most Tyr exhibiting an altered interaction network are located at the periplasmic side of BetP, Tyr550 which is in the beginning of the C-terminal domain, adjacent to the cardiolipin interaction site is another promising candidate albeit not coordinating K^+^ directly. The differences in reporter group response might also suggest an order of events triggered by cooperative K^+^ binding. We propose that the first action is a reorientation of the osmosensors from membrane bound to intratrimeric C-terminal interactions followed by a stabilization of the helical fold of the now membrane detached C-terminal domain. This two-step mechanism would involve changes in the helical fold which can be compensated at higher K^+^ concentrations.

**Figure 7.**
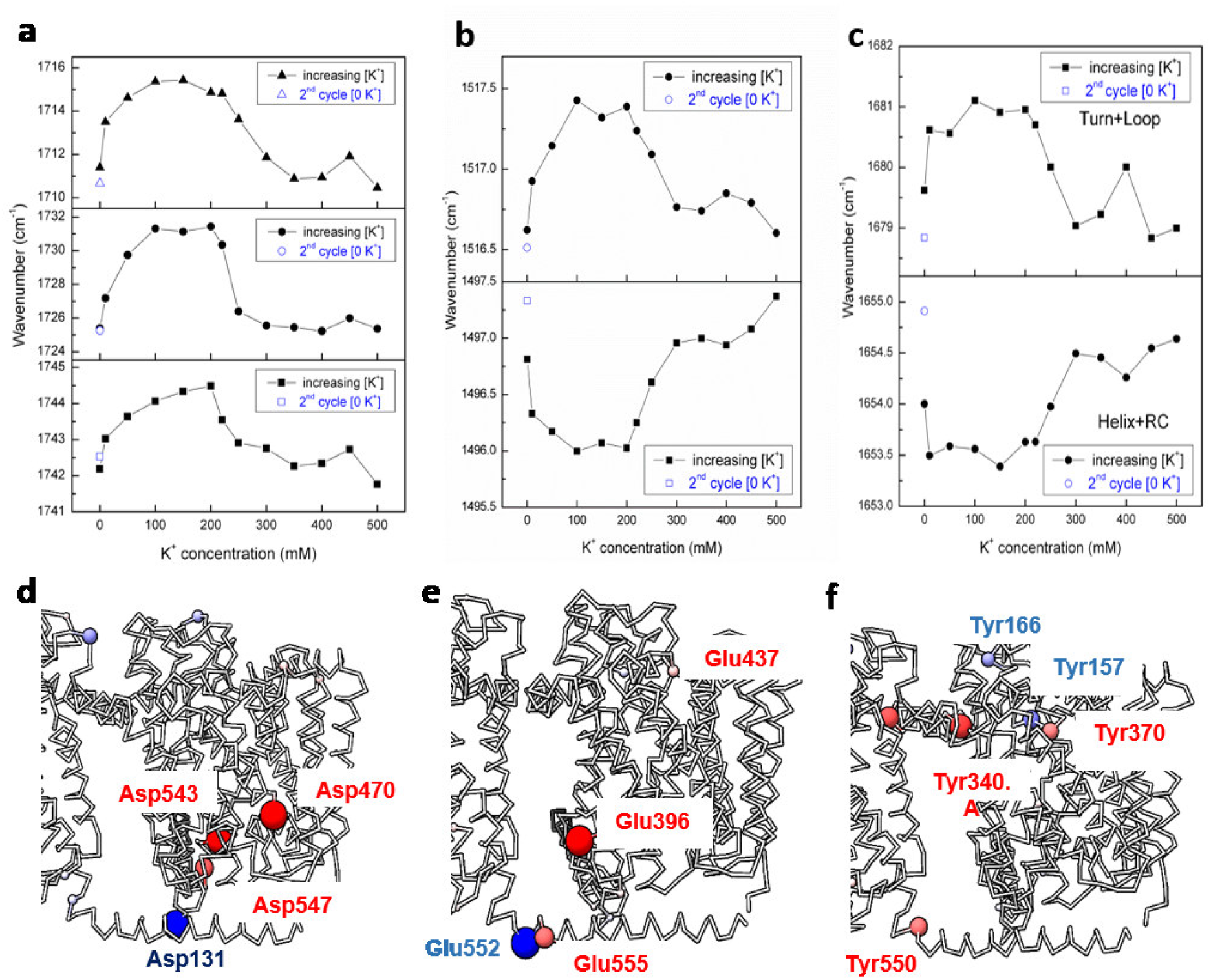
Changes in Asp / Glu / Tyr residues at different K^+^ concentrations. The shift of the peak positions with respect to increasing K^+^ concentration for **a.** protonated Asp/Glu residues absorbing at around 1712, 1726 and 1742 cm^-1^, **b.** Tyr absorbing at around 1517 cm^-1^ and 1497 cm^-1^. Open signs in blue colour represent the 2^nd^ cycle of inactive state of BetP without K^+^ ions and **c.** the secondary structures absorbing at around 1680 and 1654 cm^-1^. **d.** Aspartate and **e.** Glutamate residues, which according to the BC difference network analysis change their interaction pattern. Red spheres have stronger BC in the down-regulated state, while blue spheres have a stronger BC value in the active state. **f.** Only the cytoplasmic tyrosine residue Tyr550 showed changes from downregulated to active state. Strongest change is observed in Tyr340, which is located in h7 at the periplasmic trimerization interface. Tyr residues do not play a dominant role in the activated state except for Tyr157.

### Unfolding of the osmosensor monitored by ATR-FTIR

The IR-difference spectra recorded both in H_2_O and D_2_O also revealed indeed changes in protein secondary structure (helix, turn and loop) in response to increasing K^+^. Several peak shifts in the range of 1660 - 1630 cm^-1^ indicate a change in H-bonding strengths of ordered secondary structures both for membrane buried and solvated α-helices (**Fig. 7c**). A signal absorbing at 1653.5 cm^-1^ increases after a lag phase at 150 mM K^+^ to 1654.5 cm^-1^ and saturates at 300 mM K^+^. Reportedly, an α-helix with a high number of residues would absorb around 1653 cm^-1^ in H_2_O buffer [29], and an upshift indicates weakening or loss of H-bonding in that helix, e.g., it becomes relatively flexible. The observed increase in helix flexibility matches the increase in deprotonated Tyr residues. Simultaneously, a change in the amount of turns and loops can be assigned to the shift in wavenumbers from 1679.5 cm^-1^ to 1681 cm^-1^, exhibiting a similar hyperbolic increase as the protonated Asp/Glu residues.

To unambiguously assign the observed changes in helical structure, absorbance difference spectra of the de-regulated mutant BetP ΔC45 missing the entire osmosensory domain were recorded at 200 mM and 500 mM K^+^ (**Table S6**). The mutant exhibits minor amide I changes together with very subtle spectral alterations in the region for the protonated (above 1700 cm^-1^) and deprotonated (1600 – 1550 cm^-1^) Asp/Glu residues, as compared to WT BetP (**Fig. S14**), suggesting that these residues respond to K^+^ also in the absence of the osmosensor. Furthermore, an elevated K^+^ concentration leads to increased intermolecular β-sheet formation in the BetP ΔC45 mutant, indicated by a positive 1623 cm^-1^ signal. The secondary structural elements reveal a bi-phasic behaviour, too. While no shifts in the signal positions were detected when comparing secondary structural elements in the absence and presence of 500 mM K^+^, there is a significant difference of 2 cm^-1^ at 200 mM K^+^ in the signal positions assigned to the conformations of loops/turns and in short and 3_10_-helices. A new peak appears at 1635 cm ^-1^, both in H_2_O and D_2_O, which can be assigned to unordered structures, short helices, or solvated loops. This peak is absent in BetP ΔC45 (**Table S5**), suggesting an unfolding event of the C-terminal domain.

## Discussion

Accumulation of compatible solutes such as betaine by uptake is in the first line of defense when microorganisms face a hyperosmotic shock. Osmoregulated transporters that can sense the amount of stress and can switch accordingly from low basal transport rate to maximal rate play an essential role in immediate response. The sodium-coupled betaine symporter BetP is a prototype of such transporters, as its activity increases to more than 5-fold of the basal rate within a second [1, 30]. Biochemical data have uncovered an amazing causality between the nature of stress stimuli and transport upregulation in BetP with clear dependences on lipid environment and intracellular conditions. In fact, BetP is one of the best studied transporters for which the role in activation of practically every residue has been investigated over the past 20 years [1, 7]. In addition, multiple structures of BetP adopting individual states of the alternating access cycle describe betaine transport in molecular detail [2, 14–17]. Despite all these structural and functional data, the molecular mechanism of stress sensing and regulation in BetP remained largely unknown, mainly since crystal structures showed the osmosensor only partly resolved and pointing away from the membrane. An unambiguous assignment of the three different C-terminal conformations within one BetP trimer in a crystal structure to inactive/active states was not possible.

The combination of methods used in this study finally allows us to gain molecular insights into a complex interplay of lipids and K^+^ ions regulating BetP in response to osmotic stress. Functional data showed high cooperativity of K^+^ binding. Structural data obtained from both X-ray crystallography and CryoEM single particle analysis conclusively allowed the assignments of down- and up-regulated states and identified regulatory K^+^ interaction sites. MD simulations monitored conformational stability and validated structural states, and finally, FTIR allowed us to probe unfolding events of the terminal domains. From these data a mechanism emerges in which the C-terminal domains undergo a molecular switch to change interaction partners within the trimer. But instead of an anticipated rigid body movement, we observe a sequence of unfolding and folding events of the osmosensory helix controlled by lipids and K^+^. Concurrently, BetP protomers in the trimer re-orient themselves in the membrane thereby altering protein-protein and lipid-protein interactions to increase flexibility of helices important in transport. One major lipid interaction, which was not observed in crystal structures is with cardiolipin.

### Role of cardiolipin

The cardiolipin observed in AMP-BetP is located in a strategical position coordinated by the C-terminal domain and L2 and its presence is correlated with the orientation of the first helical segment HC1. TM12 and the C-terminal helix are connected by an elbow-like short loop around Tyr550 that needs to be flexed to position the C-terminal helix parallel to the membrane plane. Cardiolipin seems to participate in maintaining the orientation of both the elbow-loop and HC1 in the down-regulated state and enables downstream interactions with cytoplasmic loops that play a role in regulation. HC1 and HC2 folding results in the formation of the extended C-terminal helix reaching the adjacent protomer. Consequently, protomer distances are reduced at the cytoplasmic side when compared to the X-ray structures. It appears as if the C-terminal domain pulls two protomers at the cytoplasmic side closer together. Although the trimer packing is not static in MD simulations of these states, the down-regulated conformation is more compact overall. Specifically, proximity of two protomers facilitates contacts between the C-terminal domain and L6. This small loop is a crucial region for closing the cytoplasmic transport pathway in LeuT fold transporters. From our previous structural work and mutagenesis studies, we confer this loop controls the isomerization from inward to outward, which in symport is a substrate-free step and therefore relies on a low energy barrier. We speculate that the change in protomer-protomer interaction and protomer orientation at elevated K^+^ is one of the parameters involved in a cooperativity between betaine binding sites in adjacent protomers [23].

Notably, the downregulated state was not observed in crystal structures. Moreover, neither was cardiolipin, despite the observed density for a cardiolipin molecule in AMP-BetP^cryst^. We assume that additional washing steps and detergent exchange to Cymal-5 required for crystallization might be responsible for removal of cardiolipin molecules. Interestingly, HC2 in AMP-BetP^cryst^ was not folded, most likely due to missing interactions with the N-terminal domain and L6. Nevertheless, functional data have shown that even after truncation of HC2 (ΔC25) BetP is still regulated, although not to full activity. Together with the lack of a negatively charged lipid bilayer, including the missing CL, the N-terminal truncation might be one of the reasons why BetP^cryst^ always adopts an active state conformation in crystal structures. We therefore conclude that the downregulated state can be characterized by cardiolipin interacting with HC1 lying parallel to the membrane.

### The membrane stimulus

Reconstitution in AMP-nanodiscs, mimicking the negatively charged membrane belt surrounding BetP, is to date the only condition enabling studies of structural changes in the C-terminal domain. Aside of cardiolipin we identified another lipid interaction in down-regulated BetP (**Fig. 8**): POPG bound to the periplasmic side of TM2 and h7 in an extended conformation. The negatively charged, peripheral POPG like cardiolipin is in ideal positions to perceive changes in bulk lipids via lipid-lipid interactions, and to subsequently transmit the stimuli to h7. Notably, POPG and cardiolipin densities observed in the CryoEM data obtained at high K^+^ remain in contact with h7 and the C-terminal domain, respectively, thereby moving together with h7 and the C-terminal domain. In the active state POPG changes its conformation (**Fig. 8**, hyperosmotic conditions). Under hyperosmotic stress, we suggest that local changes in lipid packing trigger a movement of both lipids via their interaction with bulk lipids, thereby cause the movement of h7 and the C-terminal domain. Altered bulk lipid-lipid interactions may be the consequence of changes in curvature, lateral pressure, or lipid packing, which would directly impact the hydrophobic mismatch at the periphery of the BetP trimer. Curvature changes might result in a concerted movement of POPG and h7 causing a protomer rearrangement. Changes in both cytoplasmic cardiolipin and periplasmic POPG position might even occur simultaneously according to the amount of stress.

**Figure 8.**
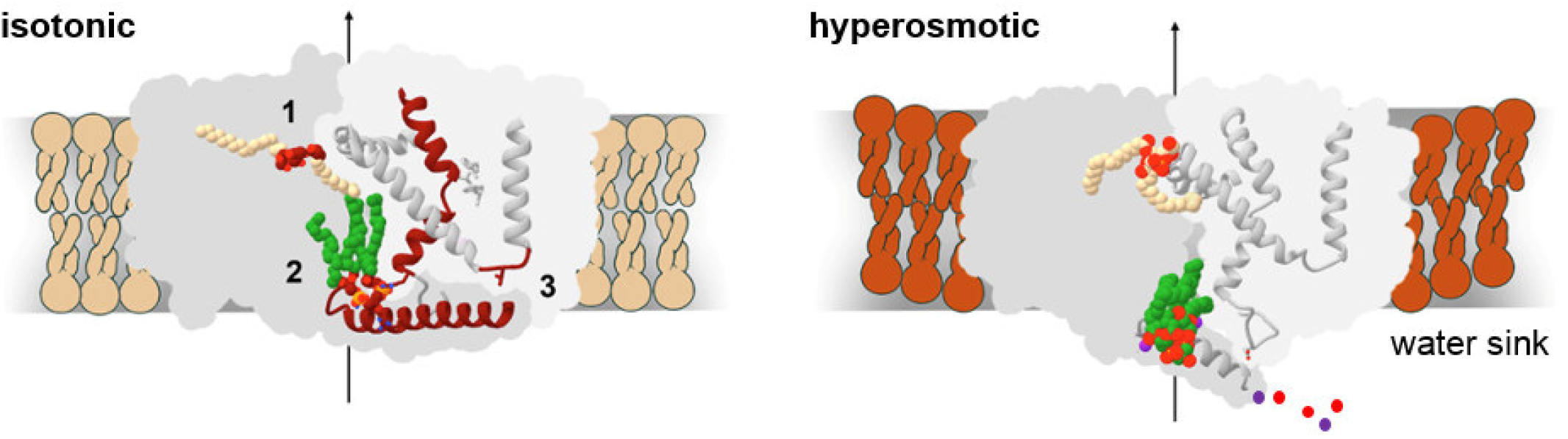
Regulation of BetP. Suggested activation mechanism of BetP. At isotonic conditions, the trimer adopts a symmetric conformation with key interactions at three positions (1, 2 and 3). Under hyperosmotic conditions, release of those three interactions leads to asymmetry in the trimer.

### How to overcome down-regulation?

The first step in activation is to overcome down-regulation when the stress signal raises above the activation threshold. For BetP, multiple interactions result in a latch-like locking of one protomer by the neighboring C-terminal domain. A comparison of K^+^ complexed CryoEM and X-ray structures suggests that the answer lies in cooperative K^+^ binding. In the CryoEM structure recorded below the *K*_d_ for K^+^, the only interaction site that is available for K^+^ binding is the one involving Tyr553 and Gln557. By contrast, in the X-ray structure, at concentrations above the *K*_d_ the C-terminal domains cross at the point where K^+^ interacts with Thr124 and Arg210. Exactly this switch in interaction – from lipid to C-terminal domain crossing – requires a flexible transition, which we suggest can only be achieved by partial unfolding of the osmosensor. FTIR shows indeed a bi-phasic behavior around the *K_d_* of K^+^ regarding protonation/deprotonation states of reporter residues and folding/unfolding events; the latter could be unambiguously assigned to the C-terminal domain by comparison to a C-terminally truncated mutant (ΔC45). This bi-phasic behavior is mirrored in betaine binding studies showing an increase at 200 mM K^+^ for sodium and betaine *K_d_* followed by a decrease at 300 mM K^+^ (**Table S1**).

### Transport activation

Interactions mediated by cardiolipin with the C-terminal domain affect the conformation of TM3 and L6 between TM6 and TM7. While in most LeuT transporters the first helix of the first repeat swings in and out, in BetP TM3a moves up and down like a mechanical spring [15]. This contraction movement in BetP is facilitated by the glycine stretch, the hallmark of betaine-specific BCCTs [31]. The glycine-stretch harbors the sodium and betaine coordinating residues Ala147, Ala148 and Met150. In the down-regulated form, TM3a is significantly retracted, with Met150 coordinating Trp374 in the betaine binding site by a non-covalent methionine-aromatic interaction. Such Met-aromatic or aromatic-Met-aromatic bridging interactions are well-established in protein structures as stabilizers [32]. A possible connection between the downregulated conformation of TM3 and the C-terminal osmosensor locking both helices of neighboring protomers might be via the beta sheet formed between L2 (125 - 127) and L8 (396 – 398) (‘2’ in **Fig. 8**, locked TM3 and the C-terminal domain are shown in red). At the same time, the interaction of the osmosensor with L6 (Lys301 - Val298) affects both L6 and TM7. L6 is stretched into an expanded position (‘3’ in **Fig. 8**), while on the periplasmic side TM7 and h7 is interacting with POPG (‘1’). Notably, TM7, is a scaffold domain helix involved in Na2 coordination via a water molecule [17]. We therefore speculate that the down-regulated state mainly affects the Na2 site, which must be occupied first to stabilize the outward facing state for subsequent betaine binding.

Under down-regulated conditions BetP might thus adopt an energetically much more stable conformation with a higher substrate affinity and a higher energy barrier to overcome during conformational cycling to the outward-facing state than the C_i_ conformation observed in the presence of K^+^ and in all crystal structures.

### The role of K^+^

As the concentration of potassium ions in the cytoplasm scales with the external osmolality, K^+^ seems like the natural choice as an activator for BetP. However, K^+^ binds with a *K*_d_ of 220 – 245 mM, a concentration already reached in *C. glutamicum* in the absence of hyperosmotic external stress, which can even reach 800 mM upon stress [18, 33]. Thus, the high potassium content is in general accompanied by high concentrations of glutamate, acting as a counter ion. Consequently, with K^+^ as physiological activator ruled out, the question remains what role K^+^ plays in activation. FTIR analysis indicates that folding/unfolding of the C-terminal domain is K^+^ responsive. K^+^ may help to transiently maintain the helical fold of the C-terminal HC1 segment, to allow for the stepwise conformational change in the transporter. The FTIR data also support a conformational change in L2 once the L6-C-terminal interaction is disrupted. Specifically, FTIR spectra for BetP-ΔC45 suggested additional K^+^ binding sites in the transporter core, where binding of K^+^ might support the isomerization from inward- to outward-facing conformation. For the C-terminal domains, we assume that K^+^ is released once the full osmosensor conformational change is achieved. Thus, we suggest a regulatory mechanism based on the symmetry-breaking unfolding/folding of the C-terminal domain affecting both the S1 betaine binding site and sodium binding sites.

Solute molecules such as K^+^ and betaine attract water molecules and restrict their freedom to move [34, 35]. An altered water potential within the cell could be an additional osmotic stress stimulus directly sensed by the C-terminal domain of BetP as water molecules are also fundamental components in protein folding by defining hydrophobic interactions. Thus, K^+^ bound to the C-terminal domain of BetP might have the sole purpose of keeping the osmosensor hydrated acting as a water sink (**Fig. 8**) at strategic positions formerly stabilized by lipid interactions. K^+^ binding in BetP is achieved by water coordination. For the folding/unfolding of BetP’s osmosensor, water potential will play a role also when internal betaine concentration increases during full activation.

### Towards a general mechanism in transporter osmoregulation

Osmotic stress induced transport regulation of BetP is considered unique even in the BCCT family as the helical folded osmosensory C-terminal domain is only conserved in *Corynebacteria* species [31]. However, anionic lipids seem to be a common denominator and are found to be crucial in other osmoregulated transporters, e.g., OpuA. OpuA is activated by increasing ionic strength, not specifically responsive to K^+^ and downregulated by cyclic-di-AMP binding to the CBS domain [13]. Despite cellular and biochemical differences in structure and transport energetics, CryoEM structures of both BetP and OpuA in downregulated and active conformations show intriguing mechanistic similarities centered around the role of negatively charged lipids. While positively charged Arg and Lys residues in BetP’s C-terminal domain - when folded - interact with the membrane to lock the downregulated conformation, OpuA harbours a comparable positively charged cluster in the helix-turn-helix motif in the NBDs, which are located close to the membrane acting as ionic strength sensor. Another striking similarity is that the CBS domains in the OpuA dimer are only ordered when bound to cyclic-di-AMP, otherwise they are natively disordered, thereby linking activation to unfolding of a regulatory domain like that reported here for BetP. In downregulated conditions, both OpuA and BetP adopt inward-facing states apparently stabilized by intra-oligomeric interactions. In OpuA, the CBS domain of one protomer interacts with the NBD of the adjacent protomer, which connects the inhibition site to the ionic strength sensor. For BetP, intradimeric interaction within the trimer, i.e., interaction between two adjacent protomers involving the cytoplasmic loops and the osmosensory C-terminal domain, is key to downregulation and appears to be facilitated by cardiolipin binding.

## Conclusions

The molecular switch model of the BetP osmosensor was predicted based on biochemical data about two decades ago; however, it could never be corroborated by structural data. Our comprehensive structural, functional, and dynamic data show that the C-terminal domain, instead of performing a rigid body movement of the entire osmosensory helix, partially unfolds. Changes in membrane curvature or lateral pressure sensed by CL-C-terminal domain interactions and/or POPG-h7 interactions may destabilize multiple cytoplasmic interactions, triggering C-terminal unfolding and disruption of interactions between the C-terminal domain and the transporter helices. Unfolding and lipid interactions of the osmosensory and regulatory domain show striking similarities to what was recently reported for the primary active transporter OpuA. In BetP, this process might be enhanced by changes in the water potential due to elevated K^+^ concentrations. The Na2 sodium ion site, and with it the energetics of cycling from inward- to outward-facing conformation, is affected by C-terminal unfolding. There are many open questions about the nature of this up-regulation, especially the role of the enigmatic Na1 site and potential K^+^ binding to the transporter itself that will require additional structural investigation in combination with studies on K^+^ dependent changes in sodium kinetics.

## Acknowledgment

This work was supported in part by the Division of Intramural Research of the NIH, National Institute of Neurological Disorders and Stroke. Anton 2 computer time was provided by the Pittsburgh Supercomputing Center (PSC) through Grant R01GM116961 from the National Institutes of Health. The Anton 2 machine at PSC was generously made available by D.E. Shaw Research. This work was supported by the Deutsche Forschungsgemeinschaft (DFG, German Research Foundation) through SFBs 699 and 1350. We want to thank the CryoEM core facility namely Bettina Böttcher are the Rudolf-Virchow-Centre at the University of Würzburg. We thank the SLS team at the PSI Villigen, Switzerland for constant support during data collection.

## Author contributions

S.M. and R.K. performed cooperativity studies. V.H., M.G.M. and C.P. determined structures, P.K and F.H. collected the Glacios data set, R.G., S.M. and O.R. performed functional experiments, G.G. and W.M. performed FTIR studies, V.L. and L.R.F. carried out MD simulations. G.G., C.P. and C.Z. designed research. C.Z., L.R.F., R.K. and V.H. wrote the manuscript.

## Material and Methods

### Cell culturing and protein purification

Cell culture and protein preparation methods have been described previously [36]. Uptake of [^14^C] betaine was measured in *E. coli MKH13* cells [37]. *E. coli* DH5αmcr [38] was used for the heterologous expression of *strep-betP*. Cells were grown at 37°C in LB medium supplemented with carbenicillin (50 μg/ml) and induction was initiated with anhydrotetracycline (200 μg/l). Cells were harvested at 4 °C by centrifugation and resuspended in buffer containing 100 mM Tris-HCl (pH 8.0) and protease inhibitor Pefabloc 0.24 mg/ml. Membranes were isolated from disrupted cells and solubilized with 1.5% β-dodecyl-maltoside (DDM) when purified protein was subsequently crystallized, or 1.0% of the same detergent when protein was reconstituted. The protein was then loaded on a StrepII-Tactin macroprep^®^ column, washed with 50 mM Tris-HCl (pH 7.5), 500 mM NaCl, 8.6% Glycerol, 0.05-0.1% DDM, and eluted with 5 mM desthiobiotin, 50 mM Tris-HCl (pH 7.5), 200 mM NaCl, 8.7% glycerol and 0.6% Cymal-5, if used for crystallization, or 0.05% DDM if used for reconstitution. Prior to crystallization the protein was loaded onto a Superose 6 (GE Healthcare) size-exclusion column equilibrated with 20 mM Tris-HCl (pH 7.5), 200 mM NaCl and 0.6% Cymal-5. This purified protein was concentrated at 4 °C to approx. 10 mg/ml at 3000 g in a Vivaspin tube (Vivascience) with a 100k -molecular-weight cut-off.

### Site-directed mutagenesis

The QuickChangeTM kit (Stratagene) and Pfu Turbo DNA polymerase were applied for nucleotide mutagenesis in pASK-IBA5betP [4] and pASK IBA7-betPΔN29-EEE44/45/46AAA [14] plasmids. All the plasmids were fully sequenced and the specific mutations confirmed.

### Crystallization and structure determination

BetP-ΔN29/E44E45E46/AAA was co-crystallized with RbCl using two different strategies. In the first strategy the purified protein was pre-incubated with Rb^+^ before crystallization trials. Therefore, purified BetP at ~8.0 mg/ml was incubated for 5 min to 16 hours in the presence of RbCl (5 – 300 mM) after which crystallization trials by the hanging drop vapor diffusion method were set. The reservoir solution, containing 100 mM Na-tri-citrate (pH 5.3 – 5.6), 100 mM NaCl, 17 – 24% PEG 400, and purified protein were mixed in 1 μl:1 μl, 1 μl:1.5 μl or 1 μl:2 μl ratios. The second implemented strategy consisted of crystallization trials using a reservoir solution in which the concentration of NaCl originally at 300 mM concentration was systematically replaced by RbCl. The concentration of betaine was varied between 1 and 8 mM. The reservoir solution was composed of 100 mM Na-tri-citrate (pH 5.3 – 5.6), 100 – 300 mM RbCl/NaCl and 17 – 24% PEG 400. Crystals grew at 18 °C and were harvested after at least two weeks. Complete datasets were collected at the tunable beamline (0.6 - 2.1 Å) PXII at the Swiss Light Source (SLS). Fluorescence scans across the rubidium absorption edge were performed to assess the incorporation of Rb^+^ and to determine the wavelength at which the anomalous scattering signal would be maximized. Data sets were collected at a wavelength of 0.81574 Å. The best data sets were collected from crystals grown in 100 mM Na-tri-citrate (pH 5.55), 300 mM RbCl and 22% PEG 400 obtained from protein without preincubation treatment. All data sets were indexed, integrated and scaled using the XDS package [39]. Processing to localize anomalous signals was performed keeping the Friedel mates separate. The quality of the anomalous signal was evaluated according to the anomalous correlation after processing in XDS [40]. The first crystal structure (PDB entry XXX) was determined by molecular replacement using the Phaser program [41], against a structure of BetP (PDB entry 3P03), whereas for the second structure (PDB entry SSS) the model of BetP in a closed conformation (PDB entry 4AIN) was used. Structures were refined using the Phenix refinement program and by manual building with the COOT program until achieving a final refined model after which any improvements of R_free_ and R_work_ factors were observed [42]. An anomalous difference Fourier map was calculated [21] with the CCP4 program FFT in order to identify Rb^+^ binding sites [43].

### Protein reconstitution into liposomes

Functional reconstitution of BetP and mutants was performed as described [36]. Briefly, liposomes (20 mg phospholipid/ml) from *E. coli* polar lipids (Avanti polar lipids) were prepared by extrusion through polycarbonate filters (100 nm pore-size) and diluted 1:4 in buffer 250 mM KPi (pH 7.5) or Tris 100 mM (pH 7.5), 500 mM KCl. After saturation with Triton X-100, the liposomes were mixed with purified protein at a lipid/protein ratio of 10:1 or 30:1 (w/w). BioBeads at ratios (w/w) of 5 (BioBeads/Triton X-100) and 10 (BioBeads/DDM) were added to remove detergent. Finally, the proteoliposomes were centrifuged and washed before being frozen in liquid nitrogen and stored at −80 °C.

### [^14^C] betaine Transport assays

Uptake of [^14^C] betaine in *E. coli* cells was performed as described [7]. *E. coli MKH13* cells expressing a particular strep-betP mutant were cultivated at 37 °C in LB medium containing carbenicillin (50 μg/ml) and induced at an OD600 of 0.5 by adding anhydrotetracycline (200 μg/l). After 2 h the cells were harvested and washed in buffer containing 25 mM KPi buffer (pH 7.5), 100 mM NaCl, then resuspended in the same buffer containing 20 mM glucose. For uptake measurements the external osmolality was adjusted with KCl. Cells were incubated for 3 minutes at 37 °C before the addition of different concentrations of [^14^C] betaine. Betaine uptake was measured at various intervals after cell samples were passed through glass fiber filters (APFF02500, Millipore, Schwalbach, Germany), and washed twice with 2.5 ml of 0.6 M KPi buffer. The radioactivity retained on the filters was quantified by liquid scintillation counting. For experiments on K^+^ stimulation in intact cells, strep-betP C252T was expressed in *C. glutamicum* strain DHPF [44], and in the same strain glycine betaine import was measured. Cultivation of *C. glutamicum* cells, manipulation and quantification of intracellular K^+^ was described previously [45]. To adjust the intracellular K^+^ concentration in *C. glutamicum*, K^+^-depleted cells were incubated for 10 min at 0.07 - 2.5 mM external KCl prior to the addition of [^14^C]-betaine. Betaine uptake assays in *C. glutamicum* DHPF were performed as described before [45] at low external osmolality of 0.24 osmol/kg to avoid osmodependent stimulation of BetP. For experiments on K^+^ stimulation in proteoliposomes, Strep-BetP C252T was purified and functionally reconstituted as described previously [36]. The internal K^+^ concentration was 72 mM, which corresponds to 40 mM KPi buffer at pH 7.5. The internal K^+^ concentration was changed by increasing the external osmolality with sorbitol, which leads to a corresponding reduction of the intraliposomal lumen. The maximal BetP activity was measured in an independent experiment at an internal K^+^ concentration of 480 mM.

### Expression, purification, and Amphipol-reconstitution of BetP

WT BetP was expressed as described previously [36]. After membrane preparation, membrane proteins were solubilized in 2% (v/v) DDM and Strep II-tagged BetP was affinity purified on a Strep Tactin® column. Reconstitution into Amphipol A8-35 was performed by mixing the polymer stock solution (100 μg/μl) with purified protein in a 1:3 ratio (w/w). After 4 h of incubation, detergent was stepwise removed by addition of BioBeads. Subsequently, the reconstituted protein sample was subjected to SEC in buffer with or without K^+^, respectively, and peak fractions were used for SPA CryoEM structure determination without further concentrating of the sample.

### Tryptophan fluorescence-binding assay

Binding assays were performed with 100 μg/ml of purified BetP either proteoliposomes, nanodiscs or amphipol A8-35. Betaine concentrations ranged from 0.05 to 15 mM. Tryptophan fluorescence emission between 315 and 370 nm was recorded on a Hitachi F-4500 fluorescence spectrophotometer and averaged over four readings, with the excitation wavelength set to 295 nm and a slit width of 2.5 or 5.0 nm for excitation or emission, respectively. The mean value and standard deviation at the 342 nm emission maximum was plotted for each substrate concentration. Binding constants were derived by fitting with the program GraphPad Prism (version 5.0c for Mac OS X, GraphPad Software Inc., San Diego CA).

### CryoEM

For SPA CryoEM structure determination, 3.5 μl of the protein sample were applied to Quantifoil R1.2/1.3 grids, blotted for 5 – 6 sec, and subsequently vitrified in a Vitrobot Mark IV (ThermoFisher Scientific, USA). Datasets were collected at a Titan Krios (ThermoFisher Scientific, USA) operated at 300 keV with a Falcon III camera (ThermoFisher Scientific, USA) in counting mode (BetP in the absence of K^+^), or at a Glacios (ThermoFisher Scientific, USA) operated at 200 keV with a Falcon III camera (ThermoFisher Scientific, USA) in counting mode (BetP in the presence of 200 mM K^+^). SPA CryoEM data processing was performed according to established procedures in RELION 3.0 and cryoSPARC V2. Symmetric and asymmetric refinements were performed for both datasets, to establish and confirm the correct symmetry, respectively. A model was built into the CryoEM density map of the down-regulated state in COOT and real-space refined in Phenix using PDB 4C7R (chain A) as a template.

### Interaction network analysis

Interaction network analysis for BetP were performed. In brief, interactions in the trimeric protein were identified in UCSF Chimera and ranked using the program Gephi 0.9.2 by applying a ForceAtlas. Ligands were excluded from the analysis. Characterization of the resulting interaction networks was performed according to the parameters ‘betweenness centrality’ (BC) and ‘modularity class’ (MC), and depicted in UCSF Chimera for visualization. Interactions were classified according to the parameter ‘betweenness centrality’ (BC). The delta-BC (**Δ**BC) values were determined to compare the C_i_^down^ *vs*. C_i_^cryst^ states, and superposition of ļ**Δ**BCļ on the C_i_^down^ structure visualizes per-residue changes in global interactions.

### FTIR

WT BetP and the C-terminal truncation mutant BetP ΔC45 were expressed and purified as described previously. For subsequent 2D crystallization, a lipid mix mimicking the lipid composition in *C. glutamicum* was solubilized into 1% Decylmaltoside (DM) at a final concentration of 4 mg/ml. Protein - lipid - detergent mixtures were adjusted to contain 2.5 mg/ml protein, 0.15% DM, and an LPR of 0.2 (w/w). After incubating overnight, the sample was dialyzed at 37 °C for 2 weeks against 500-600 ml of dialysis buffer with BioBeads added. The samples were applied to the ATR diamond, and spectra were recorded with a Vector 22 FTIR spectrometer (Bruker, Germany). Experimental details concerning data recording and analysis are provided in the supplementary methods section.

### Molecular dynamics simulations

Molecular dynamics simulations of BetP were carried out for two conformations of the full-length C-terminal domain: either that observed in the crystal structure (4C7R), or that observed in the EM map. In both cases, the rest of the protomer structure was derived from the symmetric all-inward facing crystal structure (4C7R). Each BetP trimer was embedded in a hydrated POPG bilayer, with the concentration of K^+^ ions set to either 100 mM or 300 mM. For each system, multiple independent simulations were carried out on Anton2 (with time scales from 2.4 μs to ~7.4 μs) amounting to 24 μs for the crystal structure-derived systems (12 μs for each potassium concentration tested) and 36.7 μs for the systems with the C-terminal domains in the EM orientation (19.9 μs for the 100 mM K^+^ system and 16.8 μs for the simulations with 300 mM K^+^). All simulations used the CHARMM36 force field [46–48] to describe the protein, lipid, and ions, while the water molecules were represented with TIP3P [49].

To assess whether Met150 adopts the orientation observed in crystal structures or the one observed in the EM structure, we defined two coordination numbers (*CN*_crystal_ and *CN*_EM_):

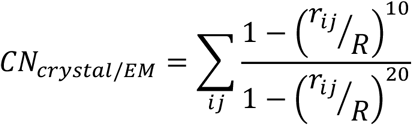

where the *i* and *j* index refers to the non-hydrogen atoms of the side chains of a given set of residues, *r_ij_* is the distance between these atoms and *R* is the distance cutoff, which was set to 3.5 Å. *CN*_crystal_ was measured using Met150, Ala313, Ala408 and Trp412 while *CN*_EM_ was measured using Met150, Trp373, Trp374 and Trp377. Consequently, if the methionine side chain is in close contact with the interaction partners observed in the crystal structure, *CN*_crystal_ will be larger, whereas if Met150 interacts with the Trp prism, then CN_u__EM_ will be larger.

