## supplementary data for "Osmotic stress response in BetP: How lipids and K^+^ team up to overcome downregulation"

**Supplementary Figures**

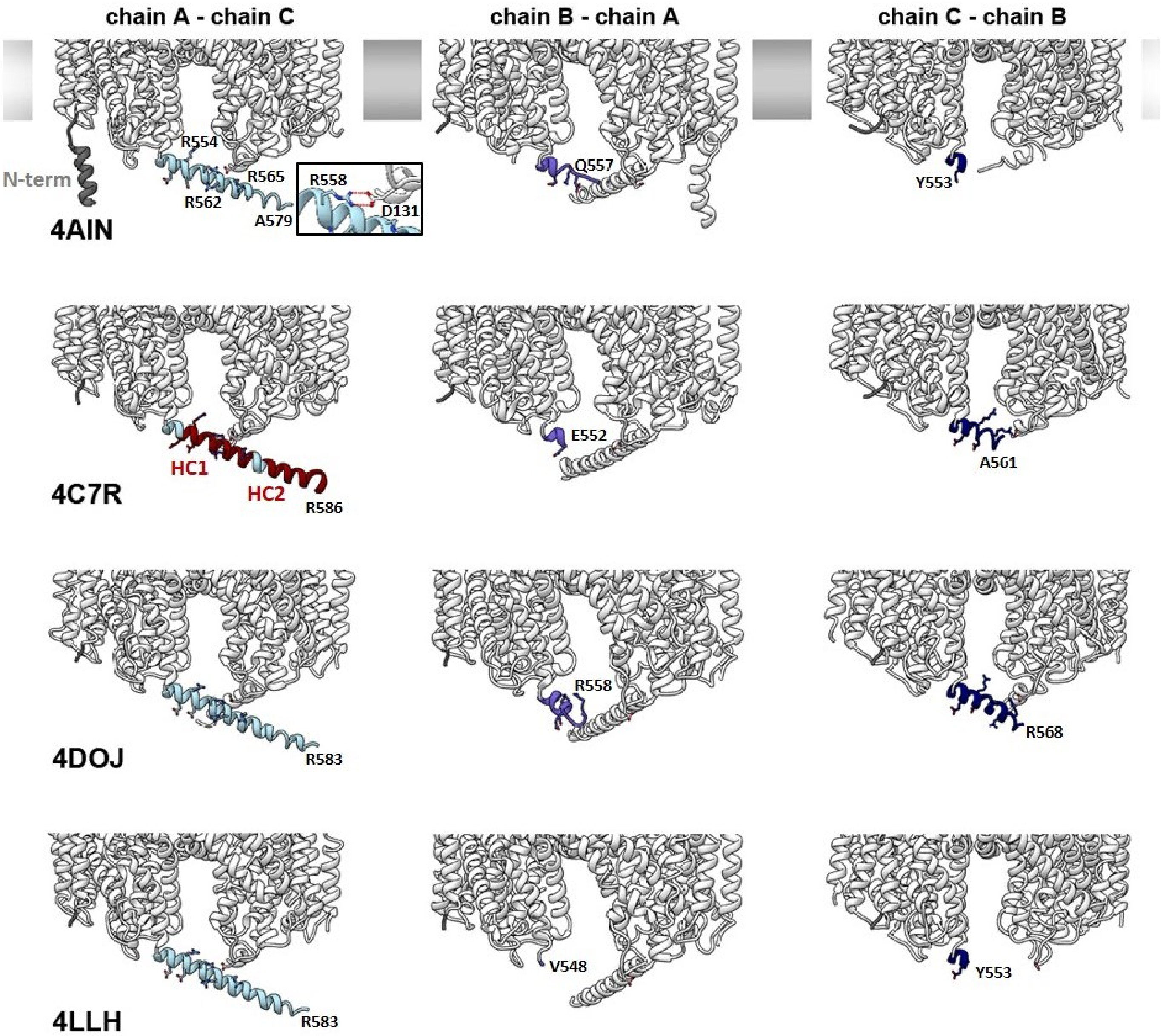


**Supplementary Figure 1.** Comparison of the C-terminal domains in BetP crystal structures
The N-terminal (grey) and C-terminal (blues) domains in BetP crystal structures 4AIN, 4C7R, 4DOJ and 4LLH exhibit differences in their resolved lengths and helical folding. In chains A, the C-terminal domains are resolved best (light blue) and the two negatively charged clusters HC1 and HC2 (marked in red in 4C7R) are helically folded. In chains B, the C-terminal domains (violet blue) are hardly resolved at all, while in chains C, they are resolved better and exhibit helical fold (navy blue). The last resolved residue is indicated for each case. The N-terminal domains (grey) are mostly unordered and not resolved in the crystal structures, except for chain A in 4AIN, where the N-terminus exhibits helical fold putatively caused by mutations introduced for crystallization. In crystal structure 4AIN, the membrane is indicated for better orientation. Strikingly, the general pattern of resolved C-terminal length and helical fold of the three protomers is consistent throughout the four structures.

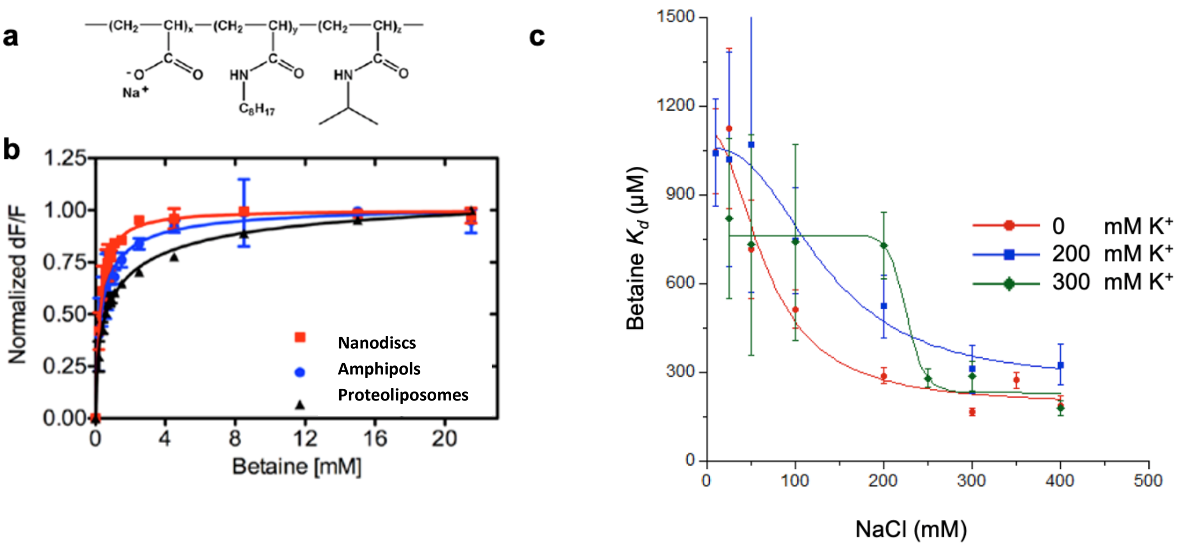


**Supplementary Figure 2.** Fluorescence absorption of BetP to show regulatory activity of BetP when **a.** reconstituted in amphipol A8-35. **b.** AMP-BetP was titrated against betaine (0-20 mM) in the presence of 300 mM NaCl and 300 mM $K^+$. Comparison of $Na^+$-coupled betaine binding to BetP reconstituted into nanodiscs, amphipols and proteoliposomes with $K_d$ values of 0.98 mM in proteoliposomes, 0.35 mM in amphipols and 0.27 mM in POPG nanodiscs. **c.** AMP-BetP was titrated at increasing NaCl concentrations against betaine (0-20 mM) in the presence of 0 mM, 200 mM, and 300 mM $K^+$. The respective betaine $K_d$ values were plotted against the NaCl concentration, and a sodium $K_d$ was determined (**Table S1**).

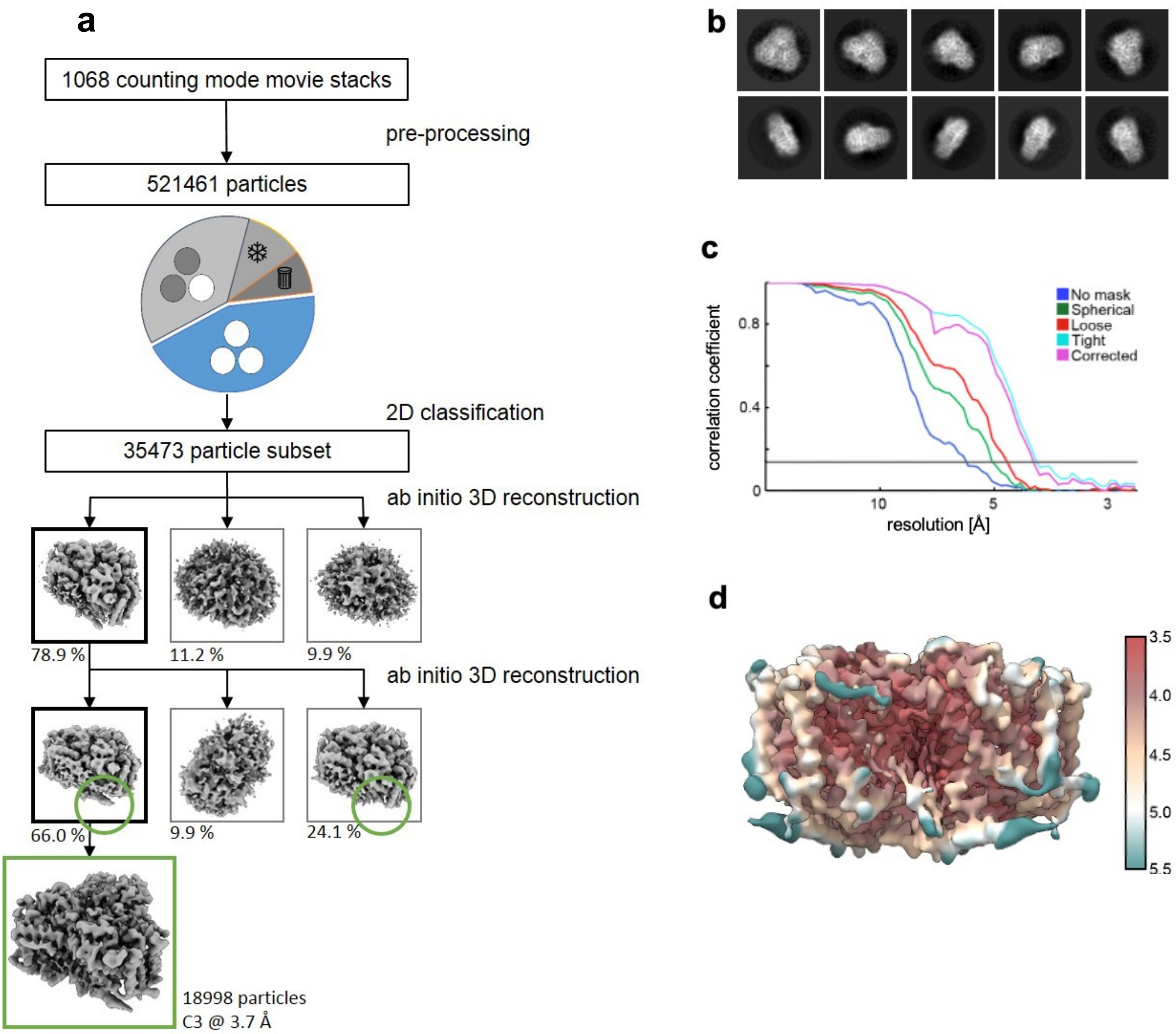


**Supplementary Figure 3**. CryoEM data processing

**a.** CryoEM data preprocessing and template-free autopicking of the Amp-WT-BetP dataset yielded a total of 521,461 particles. More than 50%, however, were identified as freezing artifacts and BetP monomers. After cleaning the trimeric BetP subset (blue fraction in the diagram) using iterative, template-based autopicking and 2D classification in RELION, 35,473 particles showing high-resolution secondary structure features were imported and processed further in cryoSPARC. Two populations were identified in the dataset. The major population (66% of particles) was attributed to the C3-symmetric, down-regulated state BetP$^{down}$, the minor population (24.1% of particles) exhibited significantly shorter C-terminal domains. **b.** Representative 2D class averages: top- and tilted views (top row) and side view (bottom row). Diameter of the circular mask is 160 Å. **c.** The FSC curve plot indicates a global resolution of 3.78 Å according to the FSC = 0.143 criterion (solid line). **d.** The local resolution distribution in the cryo-EM density map shows the rigid protein core resolved to 3.7 Å as well as more flexible parts (lipid densities, loop 6 and the last charged cluster of the sensory C-terminal domains) at a lower resolution.

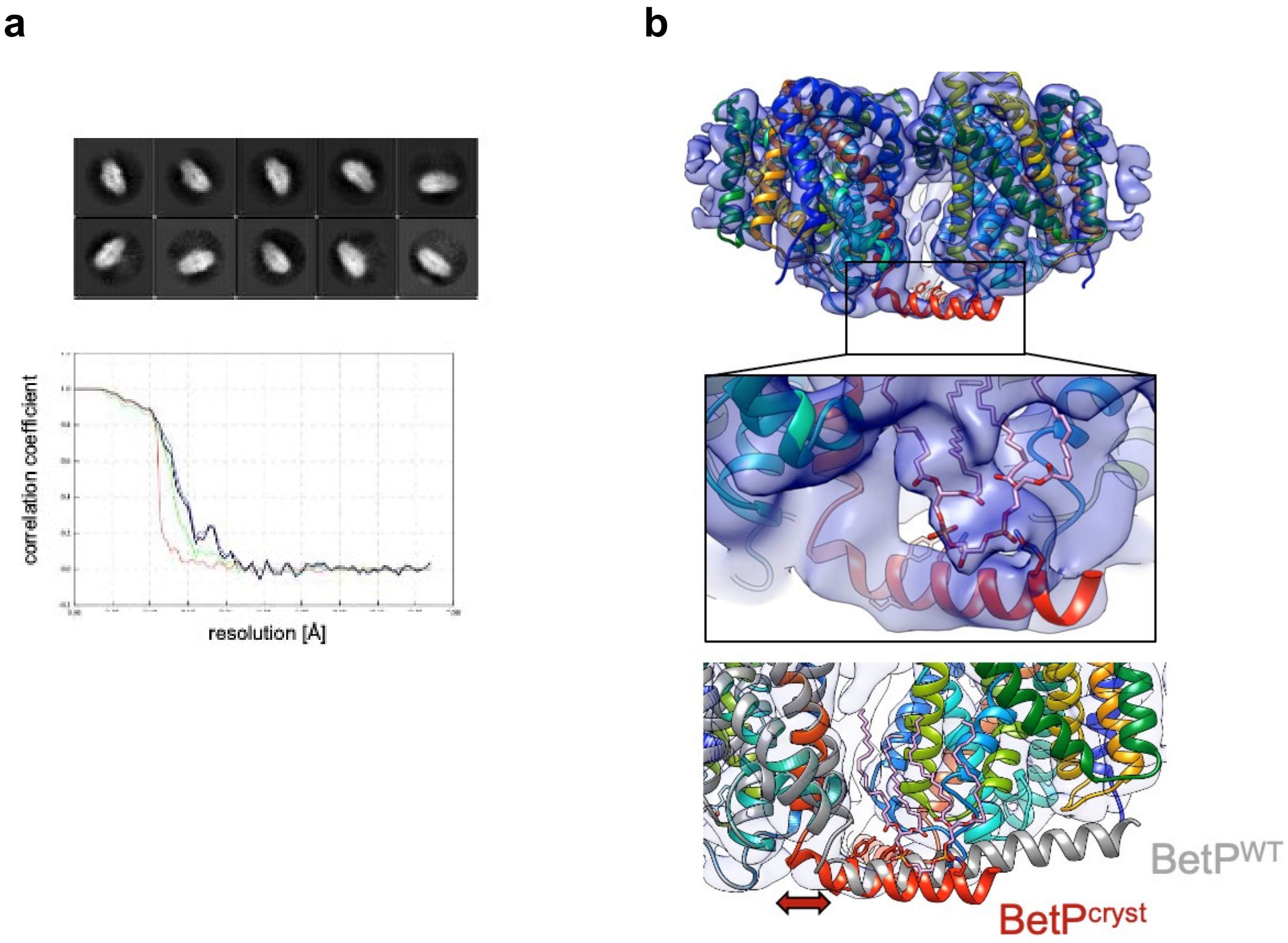


**Supplementary Figure 4. a**. Processing a CryoEM dataset of the crystallization mutant BetP$^{cryst}$ resulted in a coulomb potential map to 6.4 Å (**Table S3**). Clearly, from 2D classes side views, no density which would correspond to C-terminal domain crystal structure conformations can be observed. The data set revealed a significantly higher heterogeneity. **b.** The population distribution is shifted towards the minor population identified in Amp-WT-BetP (**Fig. S2**), resulting in the depicted coulomb potential map featuring shorter C-terminal domains with HC1 oriented still parallel to the membrane plane however slightly tilted to the outside in comparison to the WT. The elbow-loop between TM12 and the C-terminal domain is retracted towards its own protomer. A Cardiolipin density is clearly visible in AMP-BetP$^{cryst}$.

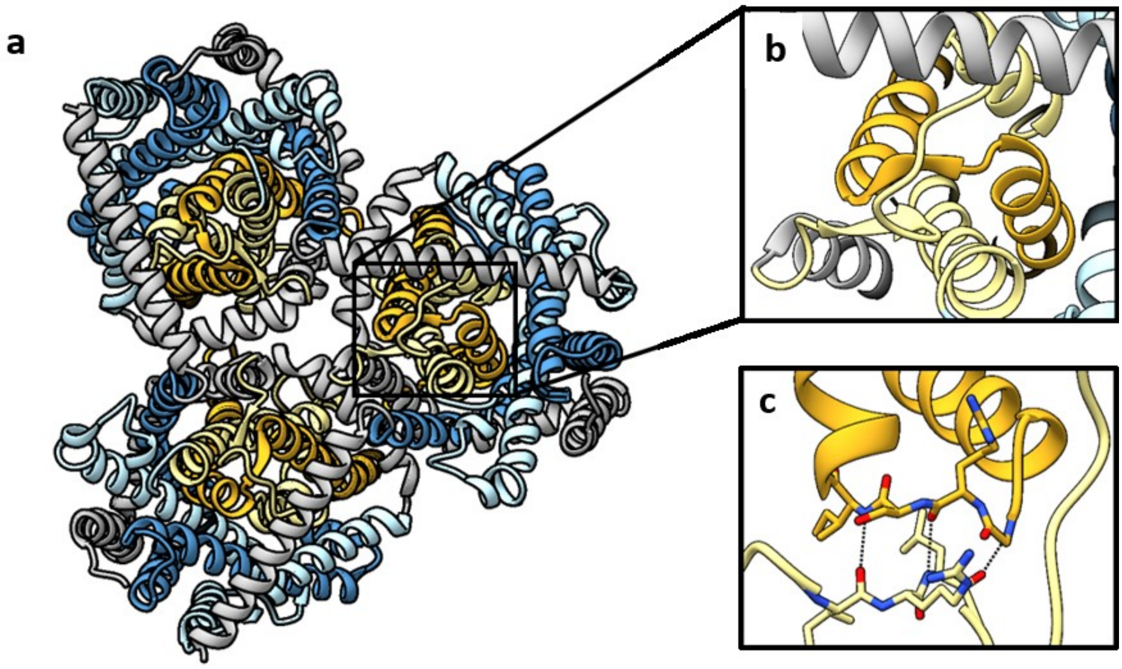


**Supplementary Figure 5.** Beta strands stabilize the cytoplasmic interaction network in the $C_i^{down}$ state.
**a.** Symmetric structure of BetP in the $C_i^{down}$ state, depicted from the cytoplasmic side.
**b.** Close up of the indicated area in a, showing a beta-sheet formed by residues Ile125 – Arg126 – Leu127 (L2) and Gly391 – Arg392 – Ser393 – Ile394 (L8). **c.** Hydrogen bonding network of the two beta strands.

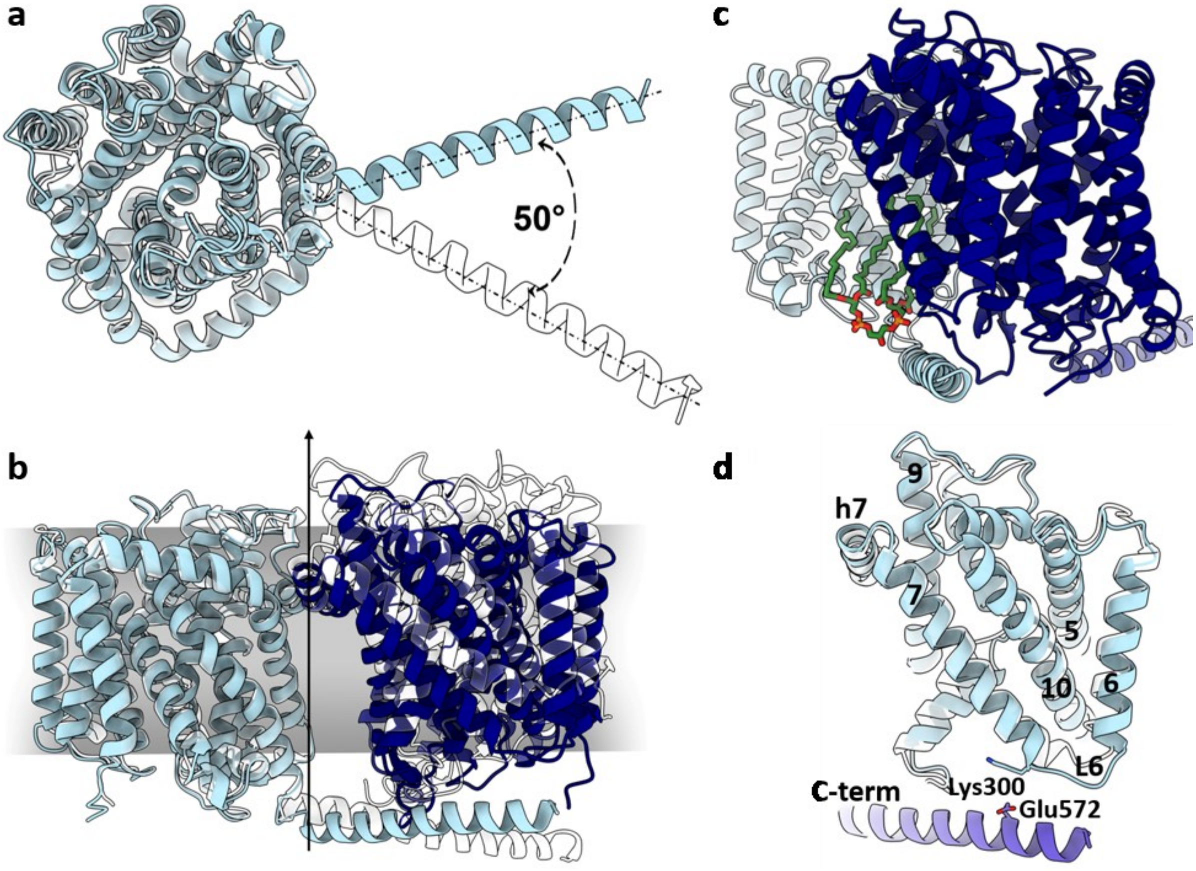


**Supplementary Figure 6.** Trimer architecture of BetP in the absence and presence of $K^+$. **a.** Superposition of protomer A in the absence of $K^+$ (light blue, cryo-EM structure) and in the presence of $K^+$ (white, crystal structure 4C7R). The orientation of the C-terminal domain varies by an angle of 50°. **b.** Superposition of protomer A (light blue) and the adjacent protomer C (navy blue) in the absence of $K^+$ (cryo-EM structure) and in the presence of $K^+$ (white, crystal structure 4C7R), aligned on protomers A. In the presence of $K^+$, protomer C undergoes an 'upward' movement with respect to the membrane normal. **c.** Interactions of the C-terminal domain in the absence of $K^+$. The osmosensor is nestled between the CL molecule (green) and L2 of the adjacent protomer. **d.** Orientation of L6 in the absence (light blue) and presence (white) of $K^+$. Without $K^+$, L6 is oriented towards the trimer periphery and the tip of the C-terminal domain, while it is facing towards the scaffold helix TM10 in the presence of $K^+$.

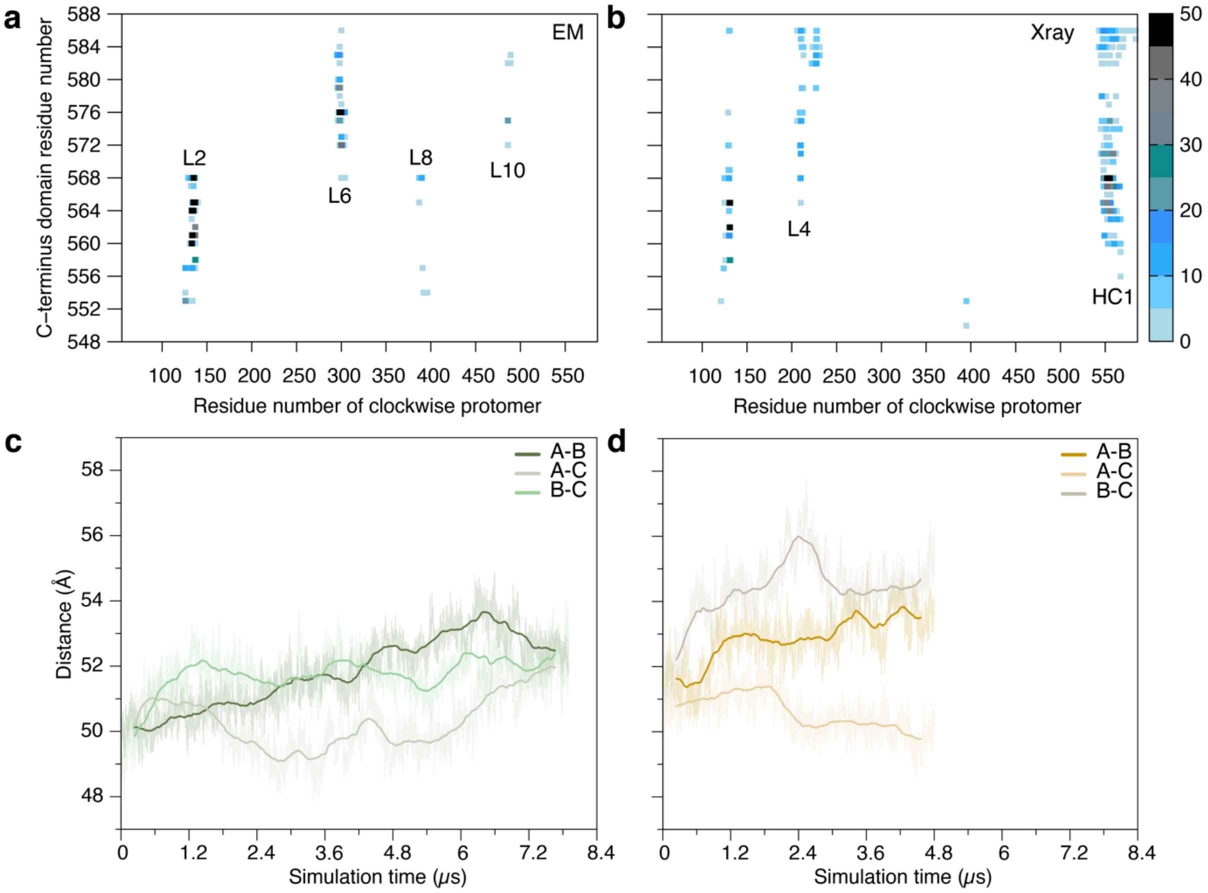


**Supplementary Figure 7.** Effect of C-terminal domain orientation on the protomer interactions during molecular dynamics simulations of BetP trimers in a POPG lipid bilayer. Pairwise contacts of the C-terminal segment with the clockwise protomer (as viewed from the cytoplasm) in **a.** the cryo-EM and **b.** the X-ray derived simulations. Contacts are defined as any two non-hydrogen atoms within 4.2 Å. Points are colored according to the frequency of each contact (%) during the simulation, averaged over the three protomers as indicated by the legend.

Centre of mass distance between the cytoplasmic segments of pairs of protomers in **c.** the cryo-EM structure and **d.** the X-ray structure derived simulation. The cytoplasmic region was defined by the residues 56-65; 113-146; 197-243; 292-307; 379-403; 471-495; 532-548. Only the atoms of the backbone were used to compute the centre of mass.

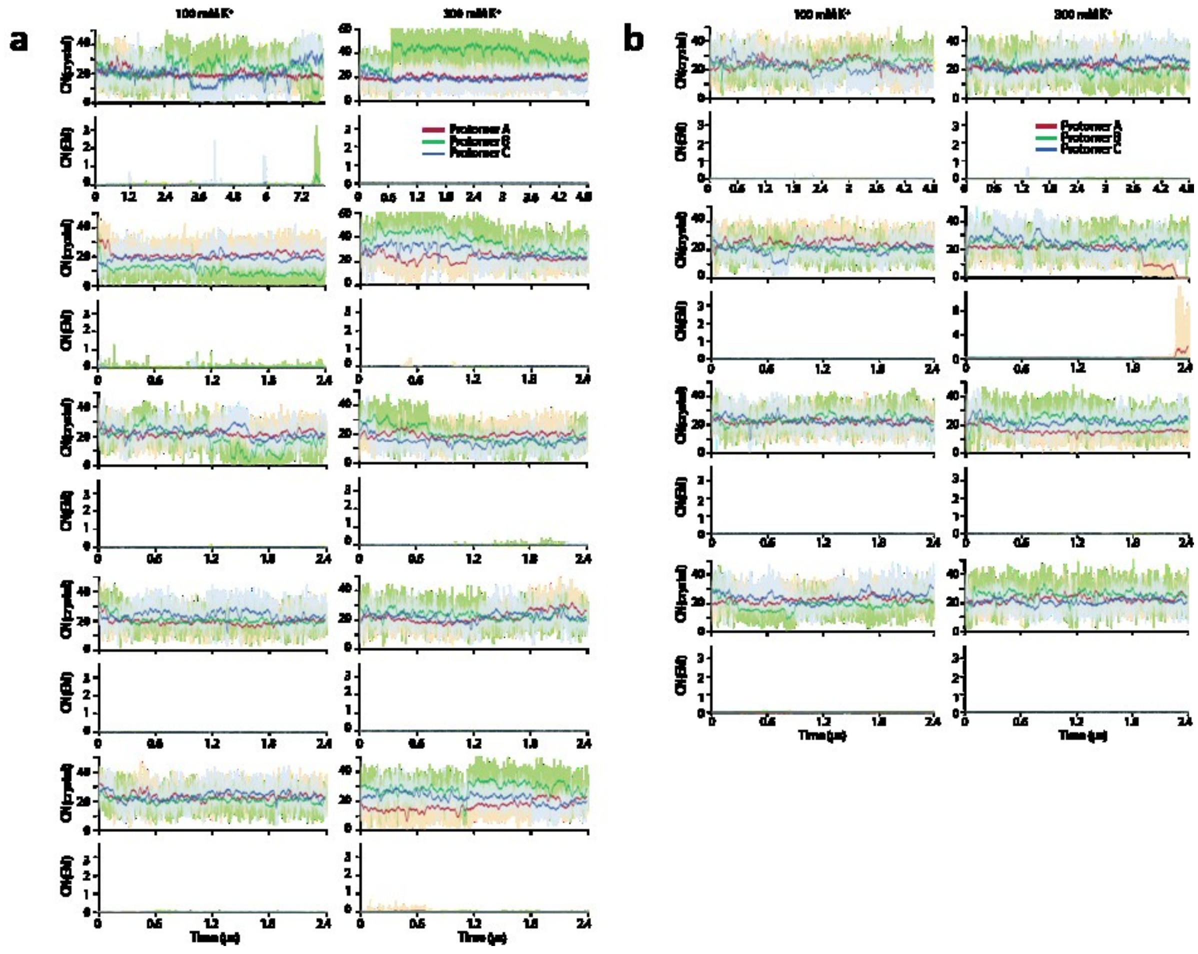


**Supplementary Figure 8**. Conformation of Met150 in multiple replica molecular dynamics simulations of the BetP trimer.

Simulations were derived from **a.** the crystal structure or **b.** the EM data. The distance of Met150 from its interaction partners is measured using a coordination number (CN) of the sidechain of Met150 with the interaction partners in the crystal structure ($CN_{crystal}$), or in the EM structure ($CN_{EM}$), as defined in the Methods. Higher numbers indicate more contacts. Simulations were carried out using (left) 100 mM $K^+$ or (right) 300 mM $K^+$.

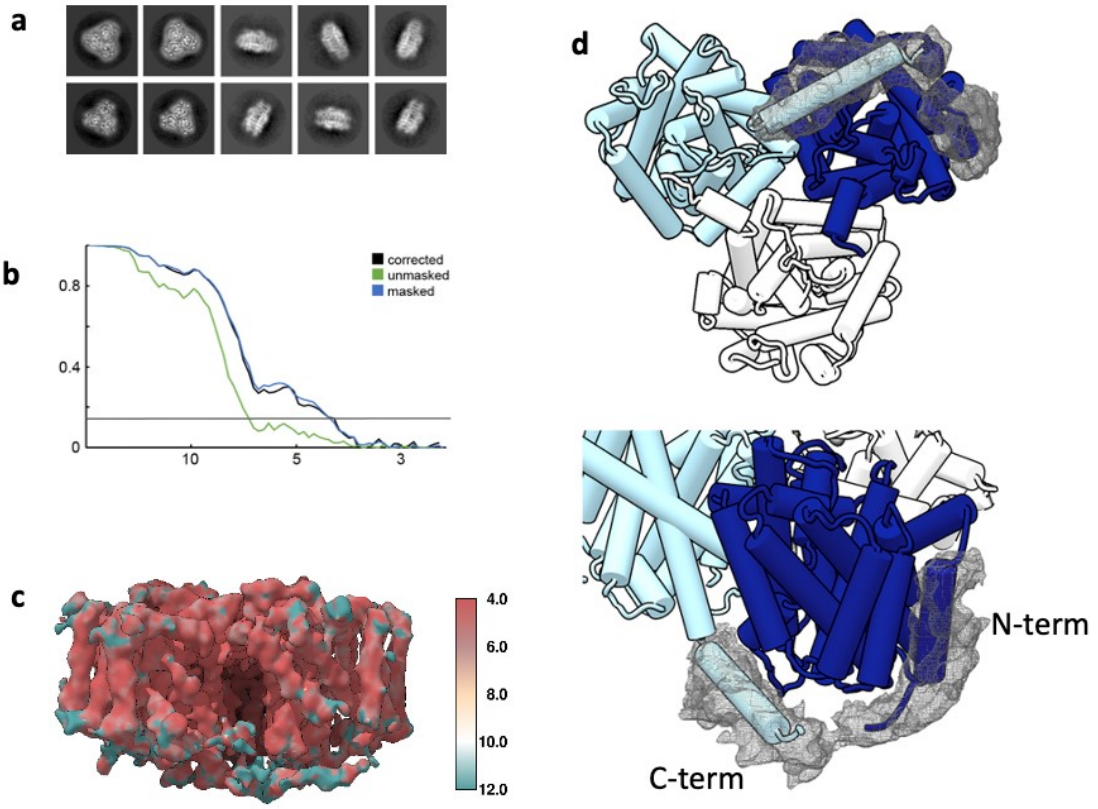


**Supplementary Figure 9**. **a.** Representative 2D class averages before (top row) and after (bottom row) Amphipol signal subtraction. Diameter of the circular mask is 170 Å. **b.** The FSC curve plot indicates a global resolution of 4.3 Å according to the FSC = 0.143 criterion. **c.** The local resolution distribution in the cryo-EM density map shows the rigid protein core resolved to 4.3 Å as well as flexible parts (lipid and sensory C-terminal domains) at lower resolution. **d.** Bottom-view and tilted view on an asymmetric reconstruction of BetP in the presence of $K^+$. The tip of the C-terminal domain in protomer A (light blue) exhibits a pronounced directionality, pointing towards TM1 of protomer C (navy blue). A weak density between the two termini (grey mesh) can be recognized when displaying the density map at an approximately twofold lower threshold (0.005) than commonly used (0.01) in UCSF Chimera.

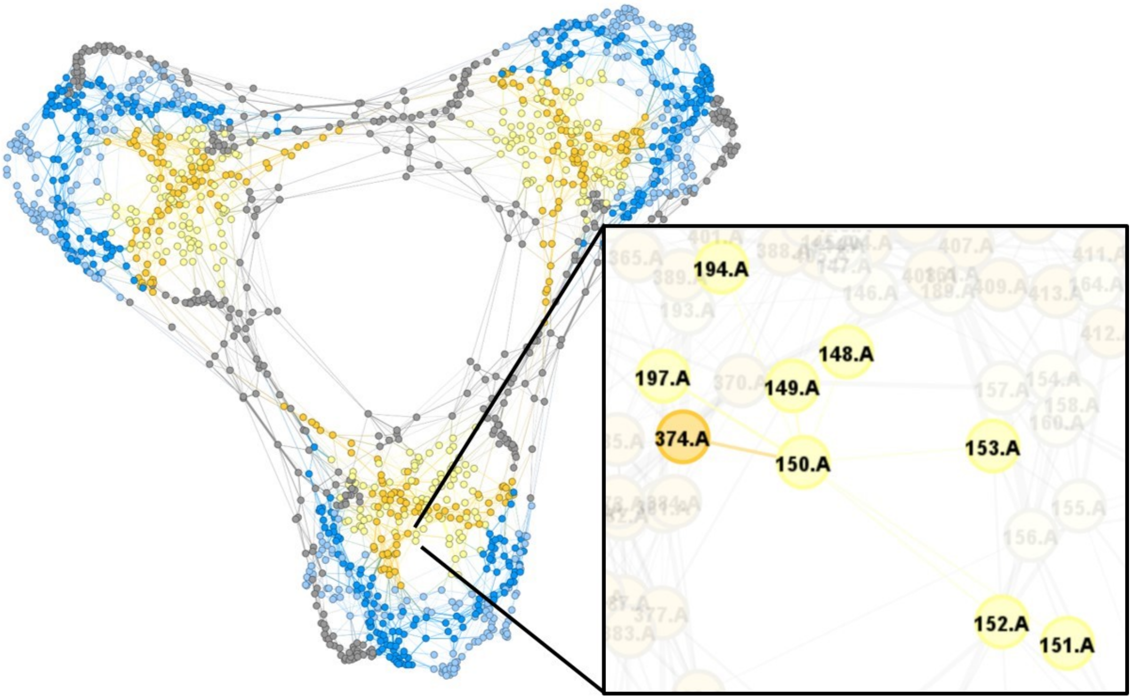


**Supplementary figure 10.** Protein interaction network of BetP in the $C_i^{down}$ state. The analysis of interactions in $C_i^{down}$ resulted in the displayed network. Protein residues are illustrated as nodes, the weighted interactions are represented by the connecting edges, color coded according to figure 4a. The zoomed window highlights the individual interaction network of Met150.A within the protein context.

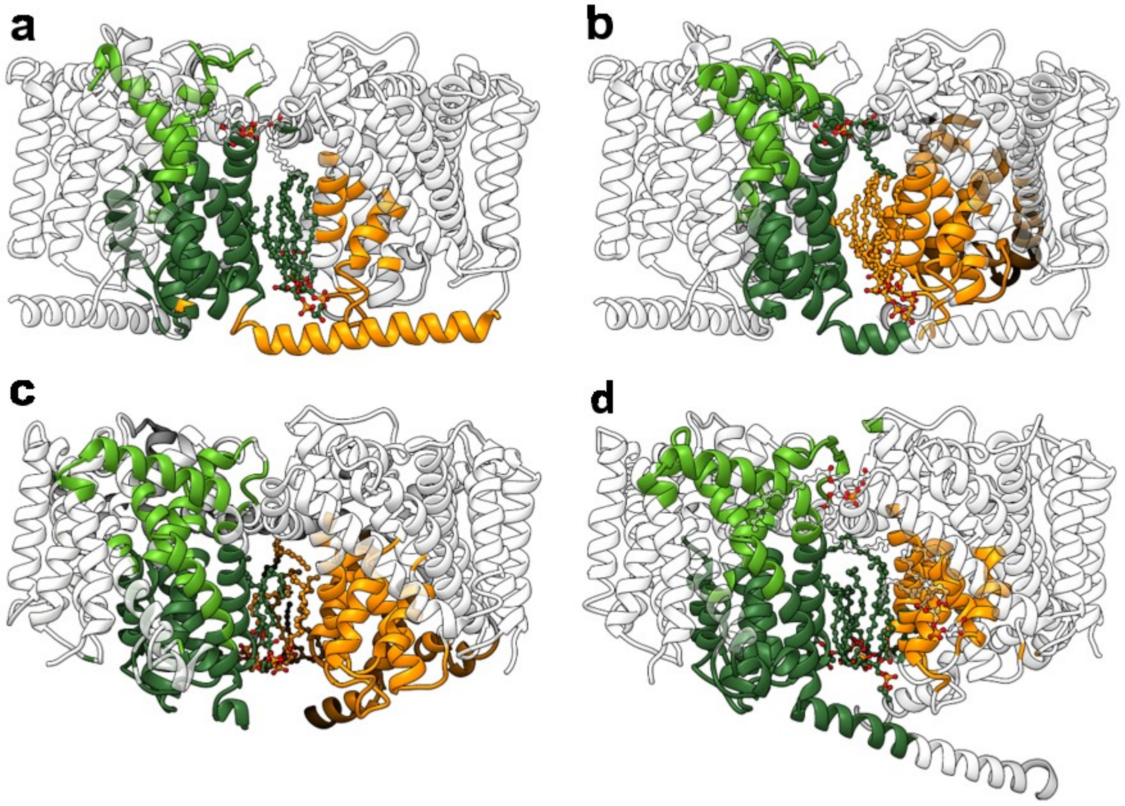


**Supplementary Figure 11.** Protein-lipid interactions of BetP during regulation.
A protein-lipid interaction network analysis was performed to determine modularity classes (MCs) for BetP in the down-regulated $C_i^{down}$ (**a**, **b**; cryo-EM structure) and up-regulated $C_i^{cryst}$ (**c**, **d**; pdb-ID: 4C7R) state. Selected modularity classes are color-coded to highlight the impact of the associated lipids on the adjacent protomers in the respective state.

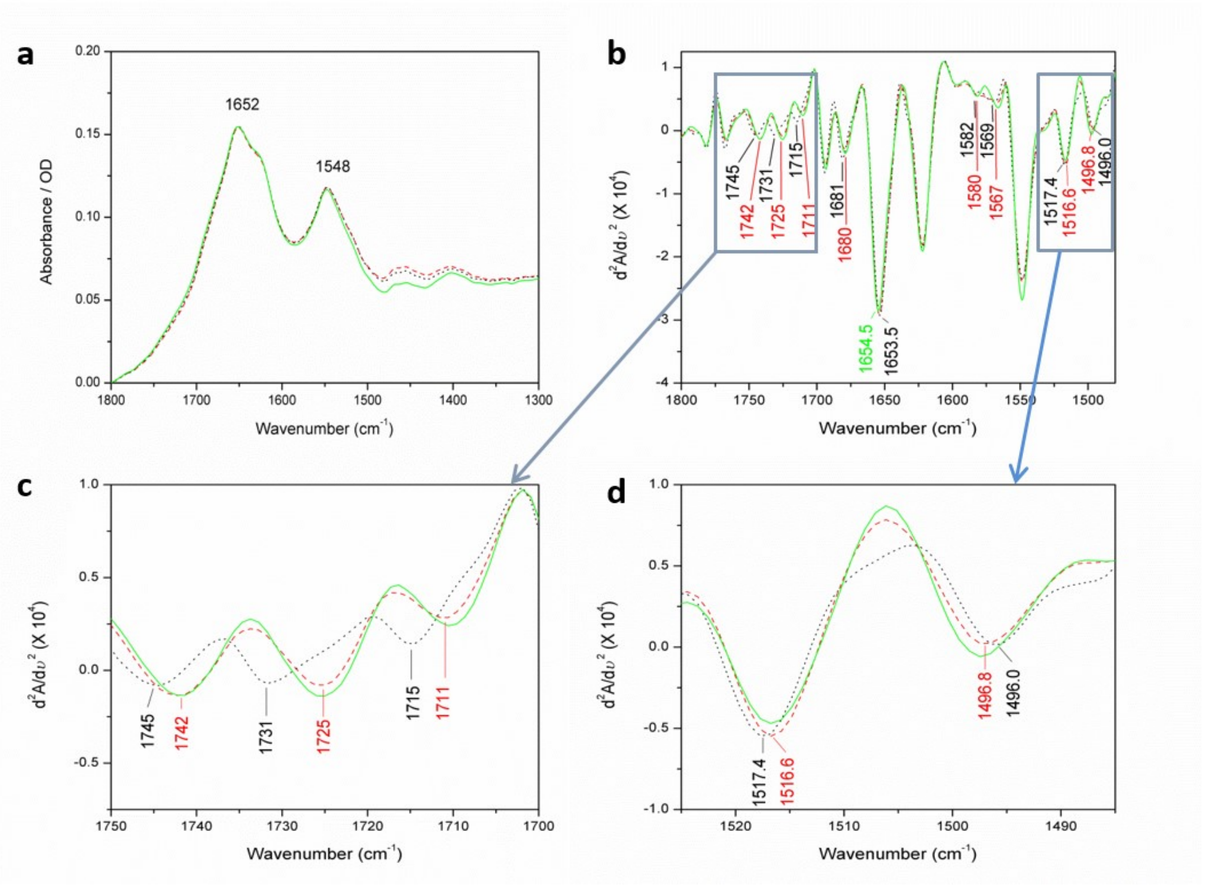


**Supplementary Figure 12.** FTIR spectroscopy analysis of BetP.
**a.** The FTIR absorbance spectra of BetP WT 2D crystals measured in the presence of 0 mM $K^+$ (- - -), 200 mM $K^+$ (….) and 500 mM $K^+$ (—). The buffer conditions were adjusted by adding various concentrations of $K^+$ to 10 mM NaPi pH 7.5, keeping the ionic strength constant. **b.** The second derivative IR spectra obtained from a. Close-up view of the spectral region marked with a box in b for **c.** the C=O modes of protonated Asp/Glu residues, and for **d.** the C-C ring stretching vibrations of Tyr residues. The labels for the peaks are colour coded.

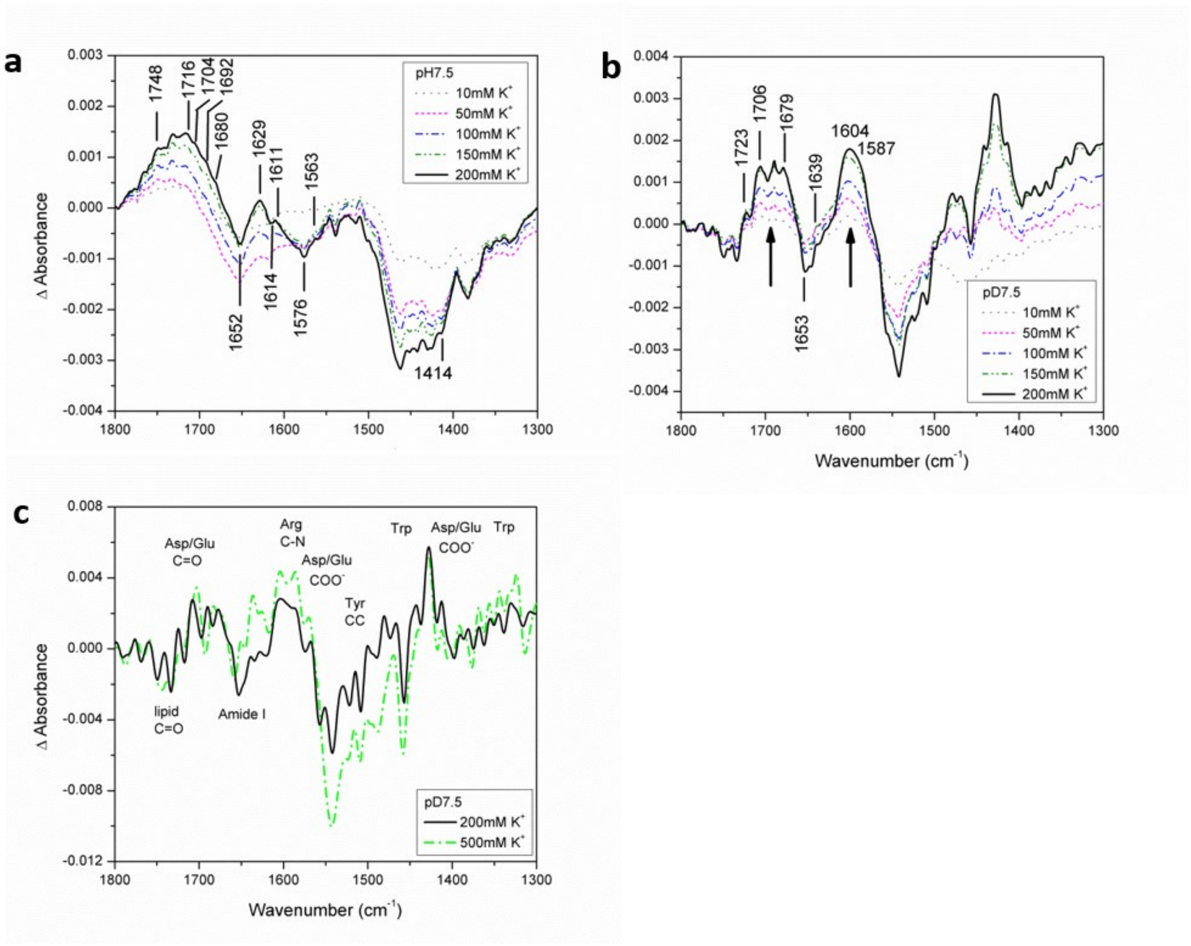


**Supplementary Figure 13.** IR absorbance difference spectra of BetP. The IR-difference spectra of BetP WT 2D crystals **a.** at pH 7.5 and **b.** at pD 7.5. All difference spectra were recorded at various concentrations of $K^+$ taking the background as inactive state of BetP without $K^+$ ions. Deuteration is required for the IR band assignments. **c.** The deconvolved difference spectrum (calculated with a Lorentzian band shape with a bandwidth of 23 $cm^{-1}$) of BetP in $D_2O$ buffer containing 200 mM $K^+$ and 500 mM $K^+$. Fourier self-deconvolution provides better resolution of overlapping peaks comprising a broad band. In deuterated buffer, the band assignment is as follows: 1745-1730 $cm^{-1}$: lipid C=O modes; 1725-1700 $cm^{-1}$: C=O stretching modes of protonated Asp/Glu; 1700-1610 $cm^{-1}$ (Amide I region): protein secondary structures like α-helices, unordered structures and loops; 1604 and 1587 $cm^{-1}$: antisymmetric and symmetric stretching C-N mode of Arg residues; 1585-1565 $cm^{-1}$ and 1390-1420 $cm^{-1}$ antisymmetric and symmetric stretching modes of deprotonated carboxylate groups for Asp/Glu, respectively; ~1517 and 1497 $cm^{-1}$: C-C ring stretching vibrations of protonated and deprotonated Tyr residues, respectively; multiple peaks at around 1482-1450 and 1350-1330 $cm^{-1}$: Trp residues.

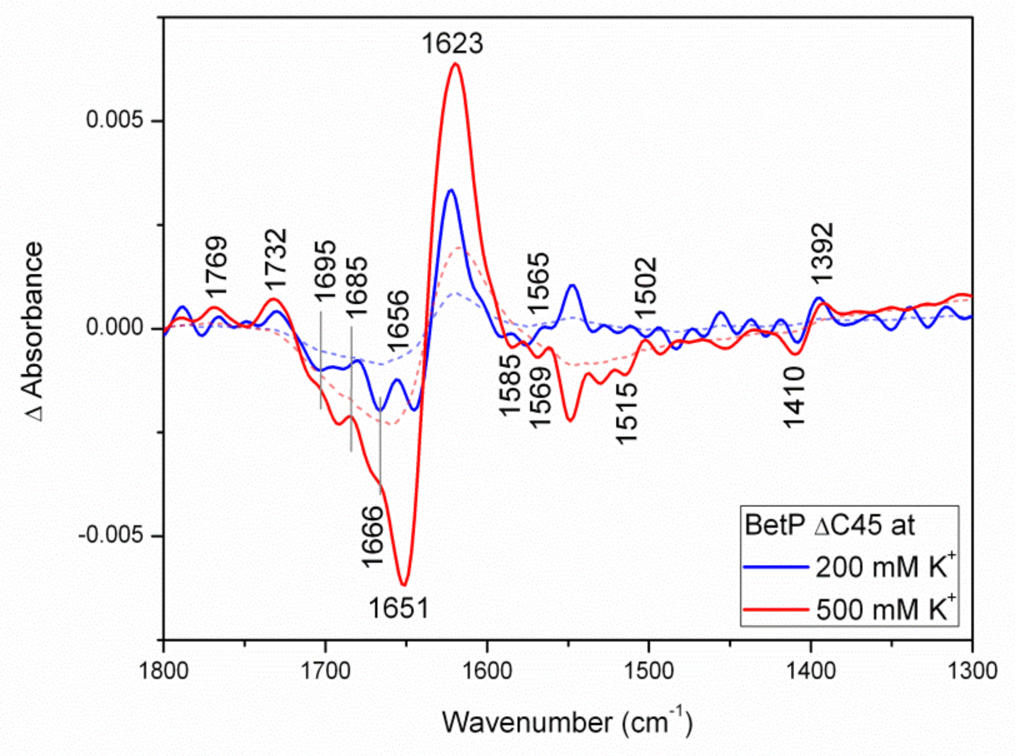


**Supplementary Figure 14.** Impact of the BetP C-terminal domain probed by IR spectroscopy. The IR difference spectra of BetP ΔC45 in 2D crystals at pH 7.5 at 200 mM and 500 mM $K^+$. Fourier self-deconvolved spectra are shown in solid lines and raw spectra in short-dashed lines. Background spectrum is recorded with perfusion of the $K^+$-free buffer (inactive state conditions).

**Supplementary Tables**

Table S1

Apparent dissociation constants $K_d$ of betaine and sodium for BetP. The values were obtained from the betaine titration evaluated by Tryptophan absorption fluorescence. Individual titration curves at 300 mM NaCl were also evaluated with respect to Hill coefficients.

| NaCl [mM] | $K_d$ [µM] 0 mM $K^+$ | $K_d$ [µM] 200 mM $K^+$ | $K_d$ [µM] 300 mM $K^+$ |
|---|---|---|---|
| 25 | 1124±143 | 821±270 | 1021±363 |
| 50 | 717±168 | 732±374 | 1071±500 |
| 100 | 514±64 | 740±333 | 747±180 |
| 200 | 288±28 | 281±30 | 524±106 |
| 300 | 188±34 | 287±49 | 351±82 |
| 400 | 167±14 | 179±24 | 326±68 |
| **$Na^+$ $K_d$ [mM]** | 66±12 | 225±13 | 113±26 |
| ***n*** | 1.3 | 1.5 | 1.9 |

Table S2
Data acquisition and processing parameters for BetP in the absence and presence of $K^+$.

| | **0 mM KCl** | **200 mM KCl** |
|---|---|---|
| **vitrification parameters** | | |
| sample buffer | 20 mM TRIS-HCl pH 7.5, 100 mM NaCl | 25 mM TRIS-HCl pH 7.5, 200 mM KCl |
| sample concentration | 1 mg/ml | ~1.5 mg/ml |
| sample volume [µl] | 3.5 | 3.5 |
| grid type | Quantifoil R1.2/1.3, Cu 400 mesh | Quantifoil R1.2/1.3, Cu 400 mesh |
| environmental conditions | 4°C / 100% humidity | 4°C / 100% humidity |
| blot time [sec] | 6 | 5.5 |
| blot force | 0 | 0 |
| **data acquisition parameters** | | |
| electron microscope | Titan Krios | Glacios |
| acceleration voltage [keV] | 300 | 200 |
| detector | Falcon III | Falcon III |
| number of movies | 1212 | 1536 |
| number of fractions | 47 | 120 |
| total electron dose [$e^-/Å^2$] | 82.21 | 30 |
| pixel size | 1.0635 | 0.9613 |
| target defocus range [µm] | -1.6 to -2.6 | -0.8 to -2.2 |
| **single particle analysis** | | |
| number of picked particles | 521,461 | 542,487 |
| particles in final reconstruction | 18,998 | 18,000 |
| particle box size [px] | 220 | 220 |
| motion correction software | MotionCor2 | MotionCor2 |
| CTF estimation software | CTFFIND 4.1 | CTFFIND 4.1 |
| 3D reconstruction software | cryoSPARC V2 | RELION 3.0 |
| applied symmetry | C3 | C1 |
| final resolution [Å] | 3.7 | 4.3 |
| resolution criterion | FSC = 0.143 | FSC = 0.143 |
| post-processing | auto-sharpening (cryoSPARC) | B-factor sharpening (B = -20) |

Table S3

Data acquisition and processing parameters for $BetP^{cryst}$ in the absence of $K^+$.

| | |
|---|---|
| **vitrification parameters** | |
| sample buffer | 20 mM TRIS-HCl pH 7.5, 100 mM NaCl |
| sample concentration | 1 mg/ml |
| sample volume [µl] | 3.5 |
| grid type | Quantifoil R1.2/1.3, Cu 400 mesh |
| environmental conditions | 4°C / 100% humidity |
| blot time [sec] | 6 |
| blot force | 0 |
| **data acquisition parameters** | |
| electron microscope | Titan Krios |
| acceleration voltage [keV] | 300 |
| detector | Falcon III |
| number of movies | 1126 |
| number of fractions | 47 |
| total electron dose [$e^-/Å^2$] | 82.5 |
| pixel size | 1.0635 |
| target defocus range [µm] | -1.6 to -2.8 |
| **single particle analysis** | |
| number of picked particles | 10,19173 |
| particles in final reconstruction | 13,234 |
| particle box size [px] | 220 |
| motion correction software | MotionCor2 |
| CTF estimation software | CTFFIND 4.1 |
| 3D reconstruction software | RELION 3.0, cryoSPARC V2 |
| applied symmetry | C3 |
| final resolution range [Å] | 6.0 to 8.0 |
| resolution criterion | FSC = 0.143 |
| post-processing | auto-sharpening (cryoSparc, Relion) |

Table S4
Model building parameters and validation for BetP in the absence of $K^+$.

| | |
|---|---|
| number of residues | 1581 |
| ligands | CDL (3), PGT (6) |
| bonds (RMSD) | |
| length [Å] | 0.005 |
| angles [°] | 0.973 |
| MolProbity score | 1.98 |
| clash score | 15.80 |
| Ramachandran plot [%] | |
| outliers | 0.00 |
| allowed | 4.00 |
| favored | 96.00 |
| rotamer outliers [%] | 0.73 |
| Cβ outliers [%] | 0.00 |
| peptide plane [%] | |
| cis proline / general | 0.00 / 0.00 |
| twisted proline / general | 0.00 / 0.00 |
| CaBLAM outliers [%] | 1.91 |
| ADP (B-factors) | |
| iso / aniso [#] | 12486 / 153 |
| min / max / mean | |
| protein | 47.72 / 174.50 / 94.26 |
| ligand | 23.05 / 116.76 / 90.32 |
| occupancy | |
| mean | 1.0 |
| occ = 1 [%] | 100.00 |
| 0 < occ < 1 [%] | 0.00 |
| occ > 1 [%] | 0.00 |
| CC (mask) | 0.78 |
| CC (box) | 0.70 |
| CC (peaks) | 0.62 |

| | |
|---|---|
| CC (volume) | 0.77 |
| mean CC ligands | 0.49 |

Table S5

| **BetP WT in $D_2O$ buffer** | | | | | | **IR Band Assignments** |
|---|---|---|---|---|---|---|
| **0 mM $K^+$** | | 200 mM $K^+$ | | 500 mM $K^+$ | | |
| **Position ($cm^{-1}$)** | Area (%) | Position ($cm^{-1}$) | Area (%) | Position ($cm^{-1}$) | Area (%) | |
| **-** | - | **1635** | 16.3 | - | - | unordered, short helix, solvated loops |
| **1640** | 11.8 | **1642** | 2.4 | **1640** | 14.0 | short helix, $3_{10}$-helix, solvated loops |
| **1654** | 74.9 | **1654** | 69.0 | **1654** | 68.4 | α-helix |
| **1665** | 7.2 | **1668** | 6.5 | **1665** | 10.8 | turns/loops |
| **-** | - | **1676** | 2.7 | **-** | - | turns/loops |
| **1679** | 6.1 | **1682** | 3.0 | **1679** | 6.9 | turns/loops |

Table S6

| BetP WT in $H_2O$ buffer | | | | BetP ΔC45 in $H_2O$ buffer | | | | **IR Band Assignments** |
|---|---|---|---|---|---|---|---|---|
| 0 mM $K^+$ | | 200 mM $K^+$ | | 200 mM $K^+$ | | 500 mM $K^+$ | | |
| Position ($cm^{-1}$) | Area (%) | Position ($cm^{-1}$) | Area (%) | Position ($cm^{-1}$) | Area (%) | Position ($cm^{-1}$) | Area (%) | |
| **-** | **-** | 1636 | 11.5 | - | - | - | - | unordered, short helix, solvated loop |
| **1645** | 46.1 | **1644** | 23.4 | **1646** | 54 | **1647** | 58.9 | α-helix + unordered |
| **1656.4** | 20.4 | **1656** | 31.9 | **1657** | 15.5 | **1657** | 10.3 | α-helix + unordered |
| **1665** | 17.9 | **1666** | 25.0 | **1664** | 22.8 | **1664** | 22.7 | unordered, short helix, solvated loops |
| **1679** | 15.6 | **1682** | 8.2 | **1679** | 7.7 | **1679** | 8.1 | turns/loops |
